# Automated Cytometric Gating with Human-Level Performance Using Bivariate Segmentation

**DOI:** 10.1101/2024.05.06.592739

**Authors:** Jiong Chen, Matei Ionita, Yanbo Feng, Yinfeng Lu, Patryk Orzechowski, Sumita Garai, Kenneth Hassinger, Jingxuan Bao, Junhao Wen, Duy Duong-Tran, Joost Wagenaar, Michelle L. McKeague, Mark M. Painter, Divij Mathew, Ajinkya Pattekar, Nuala J. Meyer, E. John Wherry, Allison R. Greenplate, Li Shen

**Author notes:** Corresponding author: Li Shen, Ph.D. - B306 Richards Building 3700 Hamilton Walk, Philadelphia, PA 19104.

## Abstract

Recent advances in cytometry technology have enabled high-throughput data collection with multiple single-cell protein expression measurements. The significant biological and technical variance between samples in cytometry has long posed a formidable challenge during the gating process, especially for the initial gates which deal with unpredictable events, such as debris and technical artifacts. Even with the same experimental machine and protocol, the target population, as well as the cell population that needs to be excluded, may vary across different measurements. To address this challenge and mitigate the labor-intensive manual gating process, we propose a deep learning framework UNITO to rigorously identify the hierarchical cytometric subpopulations. The UNITO framework transformed a cell-level classification task into an image-based semantic segmentation problem. For reproducibility purposes, the framework was applied to three independent cohorts and successfully detected initial gates that were required to identify single cellular events as well as subsequent cell gates. We validated the UNITO framework by comparing its results with previous automated methods and the consensus of at least four experienced immunologists. UNITO outperformed existing automated methods and differed from human consensus by no more than each individual human. Most critically, UNITO framework functions as a fully automated pipeline after training and does not require human hints or prior knowledge. Unlike existing multi-channel classification or clustering pipelines, UNITO can reproduce a similar contour compared to manual gating for each intermediate gating to achieve better interpretability and provide post hoc visual inspection. Beyond acting as a pioneering framework that uses image segmentation to do auto-gating, UNITO gives a fast and interpretable way to assign the cell subtype membership, and the speed of UNITO will not be impacted by the number of cells from each sample. The pre-gating and gating inference takes approximately 2 minutes for each sample using our pre-defined 9 gates system, and it can also adapt to any sequential prediction with different configurations.

## Introduction

Cytometric analysis has gained tremendous attention in immunological experiments, as a method that produces reliable, high-throughput measurements of single cells. Flow cytometry, introduced in the 1960s to separate and count immune cell subtypes, underwent a critical evolution with increased speed and parameters^1–4^. Mass cytometry, also known as cytometry by time-of-flight, was developed in 2009 to include more parameters in the analysis and avoid the difficulty of fluorescence compensation^5–7^. Although flow cytometry and mass cytometry aim for similar functionality with labeled antibodies, they do not have similar strategies to detect cell aggregates^8^. While flow cytometry uses fluorophore to label the antibodies, mass cytometry uses antibodies tagged with heavy metal ions. Each experiment in cytometry can include millions of cells and more than 20 distinct features of protein measurements. Comparing with another popular single cell profiling technology RNA-seq, cytometry can provide proteomics profile that is more relevant to clinical outcome and can detect post-translational modifications^9,10^. With such high data throughput, another advantage of cytometry over RNA-seq technology is that the cytometric experiments are better at capturing rare cell types since the overall population has largely increased. In many cases, two cell types of interest can be distinguished by the expression of a small number of proteins. Thus, the protein measurements from the cytometry experiments provide a relatively easier way to classify cell subtypes.

The most important and widely used method of subtyping cell populations is referred to as gating, which is usually a boundary or polygon defined on a bivariate density map of the entire cell population (2D density map represented by two selected protein measurements). Gating for different cell types is usually configured hierarchically^11^, to mimic the biological hierarchy of immune cell differentiation. Even though mass cytometry and flow cytometry rely on different technologies, they both employ similar gating procedures for cell type identification. However, manually defining boundaries for each cell subtype and each sample can be time-consuming and labor-intensive. Thus, algorithms have been developed that attempt to automate this task, including unsupervised clustering to identify cell subtypes^12–15^, supervised deep learning methods^16–19^, as well as some density-based gating methods^20,21^. Although unsupervised methods can target certain cell subpopulations effectively, their results often contain some remaining cell clusters that are unrecognizable and/or sometimes can fail to capture rare cell types making interpretation challenging^22^. Such methods usually lack the ability to reproduce cytometric gating from human experts, and it is hard for those methods to adapt to existing domain knowledge to satisfactorily explain the clustering results. The current supervised methods all predict on a single-cell level and thus are usually computationally expensive with large datasets. Another major challenge is that such methods typically assume that single cells have been separated from doublets, debris, and other unwanted events, a process that we call “pre-gating”. Efficient open-source methods employing pre-gating using adaptive approaches to define populations and account for variability in signal intensity between samples, however, do not exist. Indeed, current unsupervised clustering methods typically require manual “pre-gating” beforehand applying unsupervised analysis. While manual pre-gating still requires much human labor to annotate and draw polygons for the region of interest, existing bivariate pipelines also need human input to pre-define the relevant parameters. For example, those parameters include where the actual population is located as well as the approximate percentage of the target cell population. Currently, no software can configure a series of bivariate pre-gating (and gating) tasks where gates are set in a hierarchical order.

The application of deep learning has been effective in disentangling complex relationships in different cytometric domains including cell type identification and cell sorting in cytometric time series data^16–19^. Most of the existing methods focus on providing a global classification of different cell types in one step by using a multi-channel dense neural network. Moreover, these methods also usually assume pre-gated data as input, and they have not been validated for complex pre-gating tasks. In particular, one of the major challenges in performing automatic pre-gating is the technical and biological variance across different subjects. Even if the experimental protocol and the panel of measured proteins are held constant, differences in sample preparation and even instrument variation, can cause fluctuations in protein detection and precise population “shape” in the data. In addition, only predicting terminal cell types (such as naïve T cells) will prevent accurate gating of the intermediate cell types (such as T cells or lymphocytes). In the bivariate setting from manual gating, gates are configured in a tree-like structure, and at each level, the parent cell population is split into smaller subpopulations so that gating results for all intermediate cell types are obtained. The hierarchy structure guarantees the interpretability of the target cell populations. It also clearly defines some cells that are not included in the subsequent gating but still belong to the current gating step (out-of-boundary cells that are not included in any subsequent cell types). The same strategy used for manual pre-gating can be easily extended to downstream automatic gating of immune cell types, since cell-type gates have a more stable and fixed bivariate boundary compared to pre-gates. Therefore, we combined those two types of tasks and refer to all of them, onward, as “gating”.

While existing methods are suboptimal to accommodate biological variance within the protein expression data, deep neural networks with convolutional kernels have the ability to address such data challenges, using the properties of translational invariance and equivariance^23,24^. The convolutional architecture was originally designed for image classification tasks. It can detect target objects regardless of their positions in the image space and learn the general features of each target object. This versatility led to their adaptation for image segmentation tasks. The reason manual gating is challenging for automated software is that human experts have a global view of the cell density in certain protein measurement spaces so that they can visually inspect and quantify the desired cell type. Therefore, to address this challenge together with the biological and technical variance, we propose and validate UNITO, a method employing image segmentation for automated gating. UNITO converts the cytometrically derived protein expression into an image of bivariate density to enable a global identification of the cell population. Furthermore, with the ability to perform pixel-level prediction, UNITO intuitively defines the region of interest on the bivariate density maps for cytometric gating. By validating UNITO on three independent study cohorts and two cytometric modalities, we hypothesize that the framework can learn any pre-gating and gating tasks from human annotation, and then adaptively draw contours and assign labels to cells from independent data. This ability to perform inference on unseen samples without human supervision will enable applications to large-scale immunology studie s.

## Results

### Problem Statement

Flow and mass cytometry assays measure the expression of a few dozen proteins at the single-cell level, using either fluorochromes or metal ions conjugated to monoclonal antibodies as reporters. The resulting data consists of multiple channels of fluorescence or metal intensity values, which are proxies for protein expression. A typical manual analysis pipeline of this data first separates live, viable single cells from unwanted events, then gates these into different cell populations. Our goal is to automate the analysis process.

Manual gating is considered the gold standard in immunophenotyping, and the accuracy of automated methods is usually defined by comparison to a human annotator. However, this approach is sensitive to subjective choices made by one person. We consider a more robust choice of ground truth, by building consensus gates based on multiple annotators. This approach allows the estimation of disagreement between human experts, which acts like an upper bound on the performance of any analysis method. In our experiments, we show that UNITO tracks the consensus as closely as each individual annotator.

Flow and mass cytometry rely on indirect reporters for protein expression, which makes the distribution of the data susceptible to noise from unavoidable variations in sample preparation. This source of technical variation is one of the most important challenges for automated analysis. In particular, due to the high variability of channels such as DNA intercalator in the Single cell gate 1 and Single cell gate 2 process, existing automated methods usually require the data to be manually cleaned beforehand. Between different subjects, while the coordinate system for the density map is fixed, the dense population (singlets) not only moves dramatically, but the distribution also changes (**Figure 1**). The location and shape of the target cell population can differ widely between samples, which makes automatic detection over the tabular data challenging. With the UNITO framework, our goal is to create the density plot and binary mask from the protein expression matrix as input to the model and predict with an image-based segmentation method. Each gate will be trained separately, and the classifier will then be used for predicting the binary mask on independent data. The final output of the UNITO framework is the cell type label for each cell and a convex contour on the density map, which resembles the manual gates.

**Figure 1.**
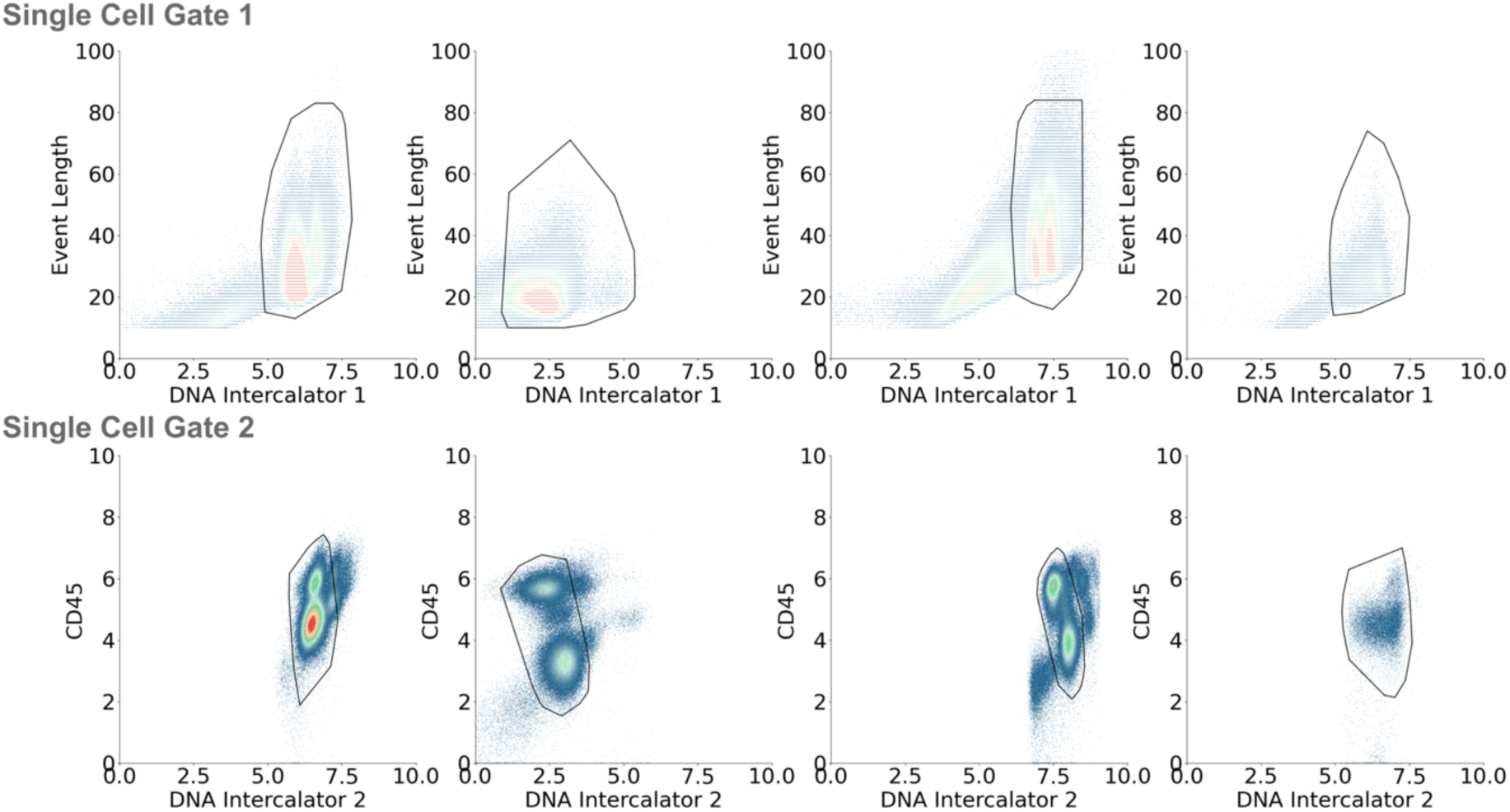
Biological and technical heterogeneity in mass cytometry across subjects. Four examples demonstrate the heterogeneity across different subjects in the first two pre-gating manual gating tasks using mass cytometry data. The top rows are single cell gate 1 and the bottom rows are single cell gate 2. The event length variable, which is the physical size of the ion cloud that results from vaporizing the cell, is an integer data, resulting in a sparser representation in the density plot. All protein markers are Arcsinh-transformed with a cofactor of 5.

### UNITO Framework Overview

UNITO uses the bivariate density plot of the entire cell population as its training data, while human annotation masks serve as the corresponding training labels (**Figure 2A**). Based on the selected measurement and label, UNITO first contains a preprocessing step that will convert and normalize the protein expression data to the density plot. Subsequently, it leverages binary labels attributed to each cell to generate an overlay mask atop the density plot. A convex hull processing will fill any empty space within the binary mask to improve the gating performance. The bivariate maps and masks are fed into the model for learning the gating pattern. The prediction output from the UNITO model is the binary mask for any independent data for validation, and the mask will undergo an additional post-processing step to interpolate the pixel label back to the single-cell classification results (**Figure 2B**). The same procedure is repeated for each gate recursively throughout the training process.

**Figure 2.**
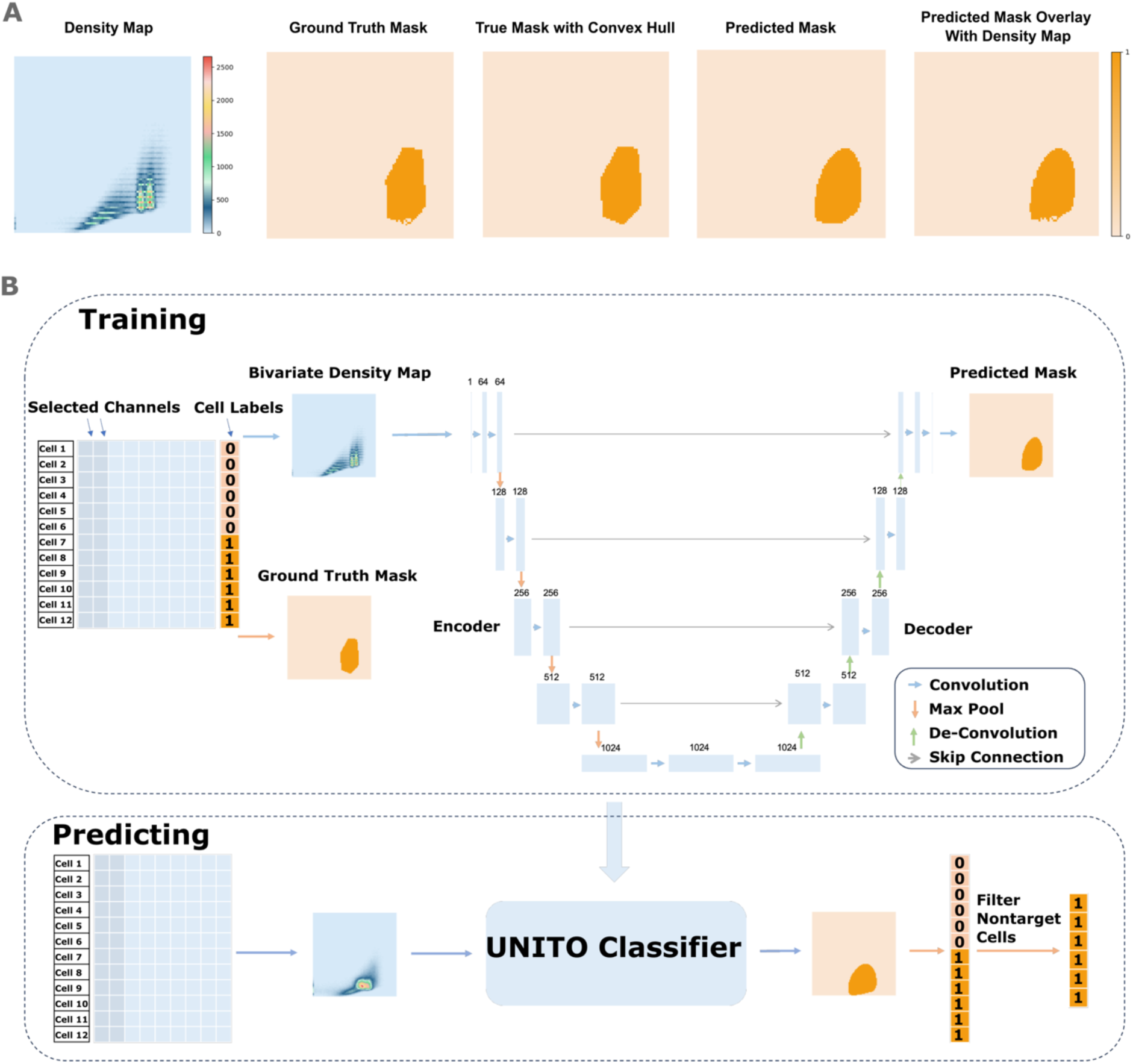
UNITO framework illustration. **(A)** Example subject input and output from single cell gate 1. The five figures from left to right are: density plot as training input, binary mask, binary mask after convex hull as training label, predicted binary mask, and reconstructed binary mask after interpolation. **(B)** Two protein measurements are selected from the expression matrix (blue matrix) and used to construct the density map, and the corresponding cell label (orange column) is used to map the overlap mask. The density maps and binary masks for the desired cell cluster are taken as training input. Prediction is performed on new density maps without annotation, and the output is the binary mask and the mapping results for all cells.

### UNITO Overall Evaluation

To validate the performance of UNITO, we constructed the ground truth gating standard by taking the consensus of multiple human annotators and compared the results with five other methods: static gates, FlowDensity^20^, FlowSOM^12^, logistic regression, and DeepCyTOF^16^. The static gates method uses the idea of density plot and binary mask construction to perform auto-gating, and every subject will get the exact same mask on the normalized space by averaging the ground truth mask. The static gating approach serves as a baseline for the proposed method. FlowDensity is a semi-automated tool built in R to gate cytometric data with positional encoding and cell population percentage. FlowSOM uses unsupervised clustering to find each cell population. Finally, logistic regression as well as DeepCyTOF use cell-level prediction by machine learning and neural network methods.

Overall, the UNITO gating for both mass cytometry data and flow cytometry data showed high correlations with the gold standard manual gating results (**Figure 3A**; **Figure S6A**). Among automated methods, the UNITO prediction was most highly correlated with the consensus gating, achieving an average correlation of 0.98 for mass cytometry and 0.97 for flow cytometry (**Table S1-2**). Moreover, UNITO had comparable, and sometimes higher, correlation coefficients than the manual gating done by individual annotators (**Figure 3B**; **Figure S6B**). To assess the performance of the UNITO framework, the average accuracy score, recall, precision, and F1 score (harmonic mean of precision and recall) were calculated across all subjects (**Table 1-2**). The accuracy score measures the number of correct predictions over all the data, the recall measures the number of true positive predictions over all of the ground true positive data, and the precision measures the number of true positive predictions over all predicted positives. Additionally, the F1 score can further disclose a more comprehensive evaluation especially when the data is imbalanced. The gating results from UNITO outperform other methods in F1 measurements for all gating tasks, and its consistency over all gating tasks guarantees its robustness for applications in flow and mass cytometric gating scenarios. In addition, UNITO uniquely performed lower-level gating tasks with high accuracy. Overall results showcased that the UNITO framework can accurately identify singlets in sequential pre-gating settings and downstream cellular populations. For its extended application in flow cytometry data, we also observed close-to-human performance that outperforms other existing methods (**Figure S6B**).

**Figure 3.**
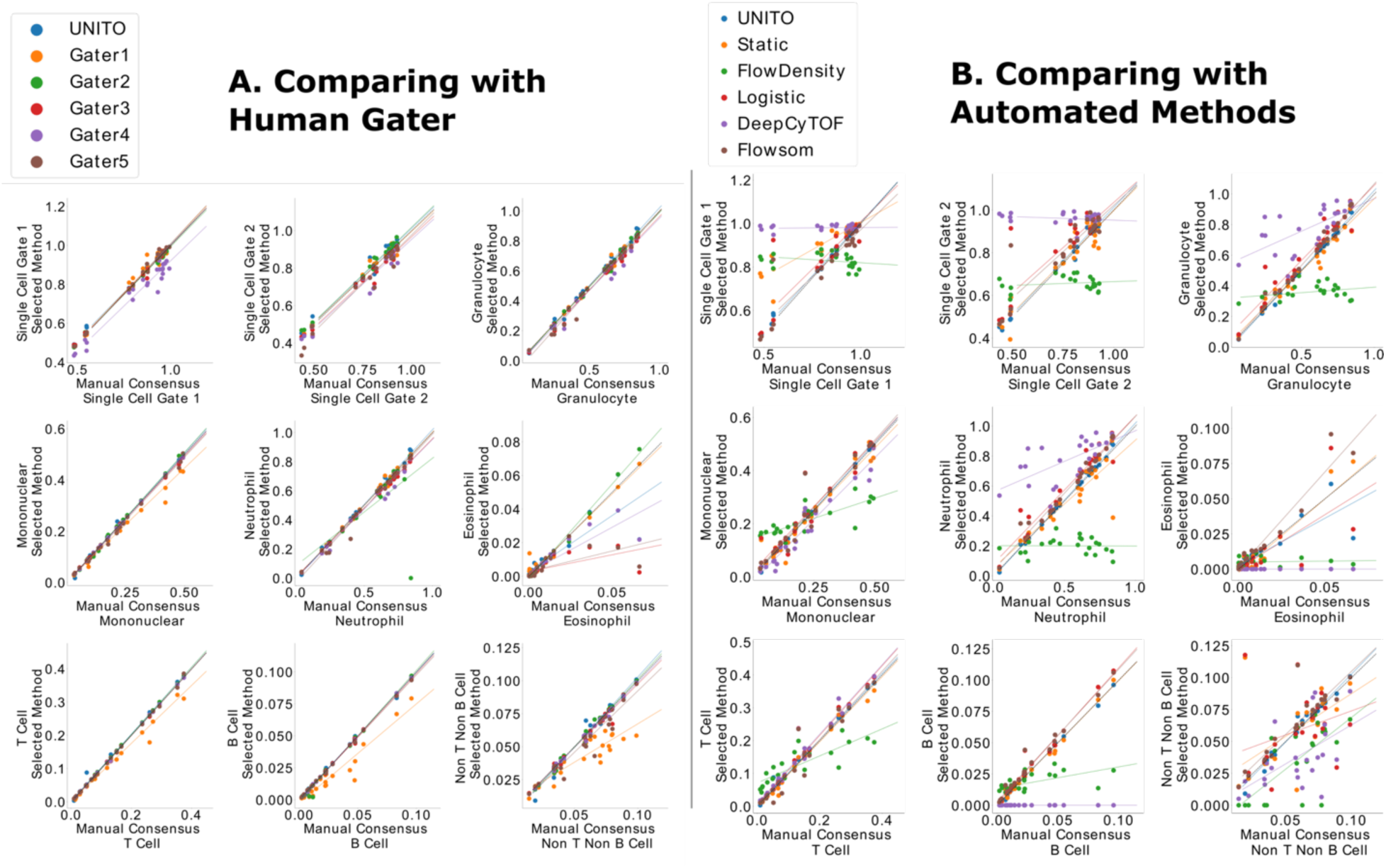
Proportion of cells over the entire cell population captured by manual gating versus human/automated gating in mass cytometry (left for vs human and right for vs automated methods). Each dot in the plot represents a single subject. The plot is separated by every single gate (including single cell gate 1, single cell gate 2, granulocyte, mononuclear, neutrophil, eosinophil, T cell, B cell, and non T non B cell), and within the same coordinate results from each method were visualized by different colors. The dashed line represents a perfect correlation with manual consensus gating, and the Pearson correlation coefficients are reported in the supplementary table S1.

**Table 1.** Evaluation matrices for cell-wise binary label classification using mass cytometry data (best scores are bolded).

|  | UNITO |  |  |  | Static Gate |  |  |  |
| --- | --- | --- | --- | --- | --- | --- | --- | --- |
|  | Accuracy | Recall | Precision | F1 | Accuracy | Recall | Precision | F1 |
| Single cell gate 1 | **0.987** | **0.997** | **0.987** | **0.992** | 0.829 | 0.892 | 0.918 | 0.859 |
| Single cell gate 2 | **0.975** | **0.993** | **0.976** | **0.984** | 0.813 | 0.823 | 0.875 | 0.827 |
| Granulocyte gate | **0.986** | **0.985** | **0.980** | **0.982** | 0.885 | 0.824 | 0.894 | 0.833 |
| Mononuclear gate | **0.990** | **0.982** | **0.975** | **0.977** | 0.932 | 0.788 | 0.840 | 0.789 |
| Neutrophil gate | **0.987** | 0.982 | **0.983** | **0.981** | 0.883 | 0.813 | 0.911 | 0.827 |
| Eosinophil gate | **0.998** | 0.863 | **0.772** | **0.790** | 0.995 | 0.632 | 0.602 | 0.516 |
| T cell gate | **0.994** | **0.984** | **0.990** | **0.973** | 0.957 | 0.785 | 0.832 | 0.792 |
| B cell gate | **0.998** | 0.972 | **0.959** | **0.963** | 0.993 | 0.779 | 0.826 | 0.780 |
| Non-T-non-B cell gate | **0.996** | 0.967 | **0.971** | **0.966** | 0.980 | 0.792 | 0.844 | 0.789 |
|  | FlowDensity |  |  |  | FlowSOM |  |  |  |
|  | Accuracy | Recall | Precision | F1 | Accuracy | Recall | Precision | F1 |
| Single cell gate 1 | 0.845 | 0.901 | 0.928 | 0.903 | 0.935 | 0.946 | 0.976 | 0.960 |
| Single cell gate 2 | 0.778 | 0.789 | 0.927 | 0.843 | 0.922 | 0.982 | 0.925 | 0.951 |
| Granulocyte gate | 0.770 | 0.672 | 0.846 | 0.728 | 0.950 | 0.967 | 0.925 | 0.944 |
| Mononuclear gate | 0.906 | 0.797 | 0.837 | 0.792 | 0.968 | 0.975 | 0.906 | 0.936 |
| Neutrophil gate | 0.722 | 0.517 | 0.880 | 0.624 | 0.956 | **0.993** | 0.919 | 0.953 |
| Eosinophil gate | 0.989 | 0.298 | 0.230 | 0.215 | 0.994 | **0.930** | 0.484 | 0.585 |
| T cell gate | 0.932 | 0.791 | 0.794 | 0.759 | 0.983 | 0.961 | 0.930 | 0.936 |
| B cell gate | 0.982 | 0.625 | 0.662 | 0.607 | 0.994 | **0.995** | 0.819 | 0.894 |
| Non-T-non-B cell gate | 0.949 | 0.472 | 0.558 | 0.503 | 0.989 | **0.979** | 0.874 | 0.921 |
|  | Logistic Regression |  |  |  | DeepCyTOF |  |  |  |
|  | Accuracy | Recall | Precision | F1 | Accuracy | Recall | Precision | F1 |
| Single cell gate 1 | 0.951 | 0.973 | 0.969 | 0.970 | 0.839 | 0.974 | 0.859 | 0.904 |
| Single cell gate 2 | 0.910 | 0.977 | 0.915 | 0.944 | 0.776 | 0.968 | 0.793 | 0.863 |
| Granulocyte gate | 0.937 | 0.985 | 0.899 | 0.938 | 0.768 | 0.971 | 0.692 | 0.789 |
| Mononuclear gate | 0.962 | 0.964 | 0.896 | 0.922 | 0.875 | 0.702 | 0.776 | 0.727 |
| Neutrophil gate | 0.938 | 0.991 | 0.893 | 0.937 | 0.758 | 0.972 | 0.677 | 0.776 |
| Eosinophil gate | 0.994 | 0.503 | 0.514 | 0.424 | 0.992 | 0 | 0 | 0 |
| T cell gate | 0.980 | 0.991 | 0.903 | 0.943 | 0.975 | 0.992 | 0.870 | 0.924 |
| B cell gate | 0.995 | 0.975 | 0.861 | 0.910 | 0.972 | 0 | 0 | 0 |
| Non-T-non-B cell gate | 0.983 | 0.878 | 0.888 | 0.863 | 0.915 | 0.279 | 0.363 | 0.289 |

**Table 2.** Evaluation matrices for cell-wise binary label classification using flow cytometry data (best scores are bolded).

|  | UNITO |  |  |  | Static Gate |  |  |  |
| --- | --- | --- | --- | --- | --- | --- | --- | --- |
|  | Accuracy | Recall | Precision | F1 | Accuracy | Recall | Precision | F1 |
| Lymphocyte gate | **0.981** | 0.996 | **0.966** | **0.981** | 0.977 | **0.999** | 0.957 | 0.977 |
| Single cell gate | **0.975** | **0.997** | 0.952 | **0.974** | 0.955 | 0.967 | 0.937 | 0.946 |
| CD3 gate | **0.988** | **0.999** | **0.966** | **0.982** | 0.963 | 0.954 | 0.930 | 0.933 |
| CD 4 gate | **0.994** | 0.989 | **0.975** | **0.977** | 0.980 | 0.955 | 0.946 | 0.939 |
| CD 4 naïve gate | **0.991** | **0.986** | **0.924** | **0.945** | 0.982 | 0.950 | 0.877 | 0.897 |
| CD 8 gate | **0.994** | **0.998** | **0.944** | **0.968** | 0.989 | 0.944 | 0.912 | 0.919 |
| CD 8 naïve gate | **0.997** | **0.992** | **0.986** | **0.935** | 0.993 | 0.933 | 0.759 | 0.803 |
|  | FlowDensity |  |  |  | FlowSOM |  |  |  |
|  | Accuracy | Recall | Precision | F1 | Accuracy | Recall | Precision | F1 |
| Lymphocyte gate | 0.695 | 0.900 | 0.650 | 0.743 | 0.927 | 0.900 | 0.950 | 0.924 |
| Single cell gate | 0.800 | 0.799 | 0.810 | 0.791 | 0.937 | 0.905 | **0.962** | 0.932 |
| CD3 gate | 0.881 | 0.748 | 0.920 | 0.810 | 0.980 | 0.978 | 0.961 | 0.969 |
| CD 4 gate | 0.907 | 0.744 | 0.870 | 0.777 | 0.989 | **0.992** | 0.956 | 0.973 |
| CD 4 naïve gate | 0.899 | 0.791 | 0.521 | 0.586 | 0.975 | 0.981 | 0.812 | 0.884 |
| CD 8 gate | 0.774 | 0.445 | 0.181 | 0.246 | 0.984 | 0.983 | 0.847 | 0.908 |
| CD 8 naïve gate | 0.825 | 0.523 | 0.066 | 0.109 | 0.996 | 0.913 | 0.870 | 0.887 |
|  | Logistic Regression |  |  |  | DeepCyTOF |  |  |  |
|  | Accuracy | Recall | Precision | F1 | Accuracy | Recall | Precision | F1 |
| Lymphocyte gate | 0.897 | 0.868 | 0.917 | 0.890 | 0.173 | 0.280 | 0.228 | 0.240 |
| Single cell gate | 0.903 | 0.879 | 0.915 | 0.895 | 0.264 | 0.211 | 0.237 | 0.205 |
| CD3 gate | 0.959 | 0.936 | 0.941 | 0.938 | 0.656 | 0.095 | 0.080 | 0.086 |
| CD 4 gate | 0.981 | 0.976 | 0.937 | 0.956 | 0.767 | 0.100 | 0.071 | 0.082 |
| CD 4 naïve gate | 0.984 | 0.953 | 0.899 | 0.923 | 0.880 | 0.097 | 0.068 | 0.078 |
| CD 8 gate | 0.986 | 0.928 | 0.909 | 0.917 | 0.909 | 0.032 | 0.038 | 0.033 |
| CD 8 naïve gate | 0.996 | 0.925 | 0.859 | 0.886 | 0.977 | 0.006 | 0.023 | 0.008 |

### UNITO Gating on Mass Cytometry Data

We next asked whether UNITO’s density map with decision boundary for the sequential gating task visually aligned with the contour from human experts’ consensus (**Figure 4A**; **Figure 2**). The ground truth boundary derived from consensus gating is almost the same as the predicted contour in all gates. Since different gates within the same gating hierarchy have the same coordinate space, those gates are plotted together. The first two pre-gating steps in the sequential prediction task are more complex than cell subpopulation gating tasks, because the presence of debris cells and doublet cells for both gates usually differ in its distribution and position in the density plot across samples. This challenge, if not manually cleaned, will largely impact the subsequent gating results. The gating output from UNITO also showed a high correlation between the proportion of predicted cells with manually gated cells as a nearly straight line in combination with all gating results from UNITO (**Figure 4B**), indicating its high consistency with the consensus gating results and ability to perform gating tasks at a level similar to human.

**Figure 4.**
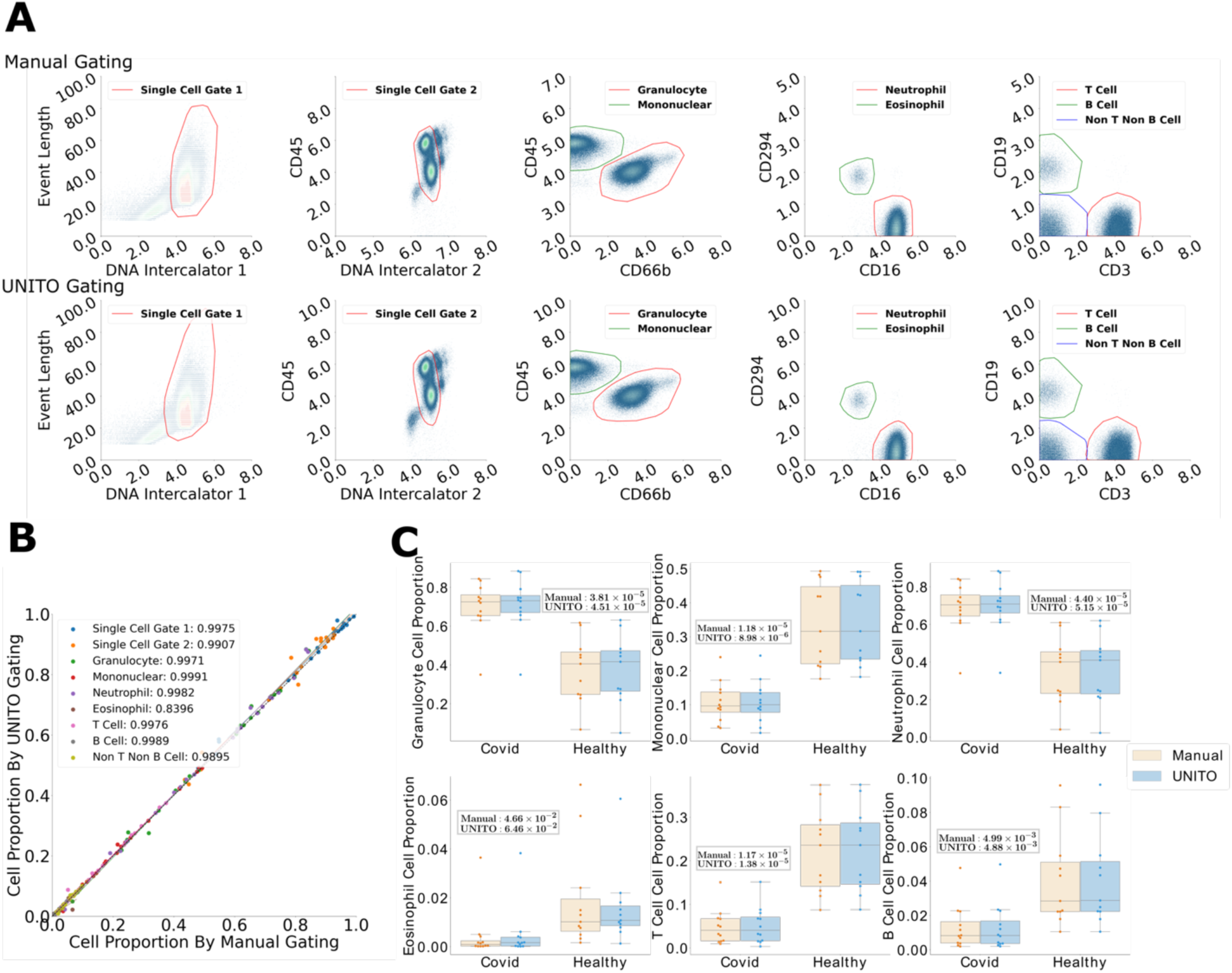
Comparison between UNITO automated gating and manual consensus gating (mass cytometry). **(A)** Density map and target cell annotation for mass cytometry data. The first row shows the ground truth annotation for one validation subject in the sequential prediction task, and the second row shows the predicted decision boundary from the reconstruction of the UNITO output. Each column represents a single level in the gating hierarchy, and gates within the same level are plotted together, such as granulocyte and mononuclear. **(B)** Visualization of the proportion of cells from UNITO and manual consensus gating with all gating tasks together. **(C)** Statistical comparison validated on the COVID-19 vs Healthy mass cytometry data (Top row: Granulocyte, Lymphocyte, Neutrophil. Second row: Eosinophil, T cell, B cell). The t-test between COVID-19 and Healthy group was applied on selected gates and P-values were reported for both manual gating and UNITO gating. For each gating, the plot is separated by subjects’ health condition while the results from manual gating and UNITO for the same population are put next to each other for easier observation.

Another key feature of manual annotation is the ability of the expert to identify cells not only by whether the cells that fall into a gate, but also if they are excluded by a gate. UNITO framework mimics this feature by excluding out-of-gate cells in gate 1 as input for the next prediction gate. This pre-filtering step can also be visualized as the difference in the entire cell density plot between manual gating and automatic gating. The decision boundary from UNITO provides a convex hull that is similar to the human annotation process, which can offer post-prediction adjustment based on the vertex of the convex hull (**Figure 5A**). If we visualize using cell-level prediction results, it will not make sense in the bivariate visualization. For image and density-based estimation, all the cells within the boundaries are classified as the target cell type, but when we visualize the results from previously published cell-level prediction, not all cells within the gates are classified by the methods (e.g. FlowDensity), or a simple convex hull over all labeled data may also include cells that are not classified as the target population (e.g. DeepCyTOF). This becomes more apparent when we show the binary distribution of all the cells classified as the target cell type in the same coordinate space as the density map (**Figure 5B**). We can see that for both convex hull visualization and binary mask, the UNITO’s prediction is the closest estimation approaching the manual consensus gating.

**Figure 5.**
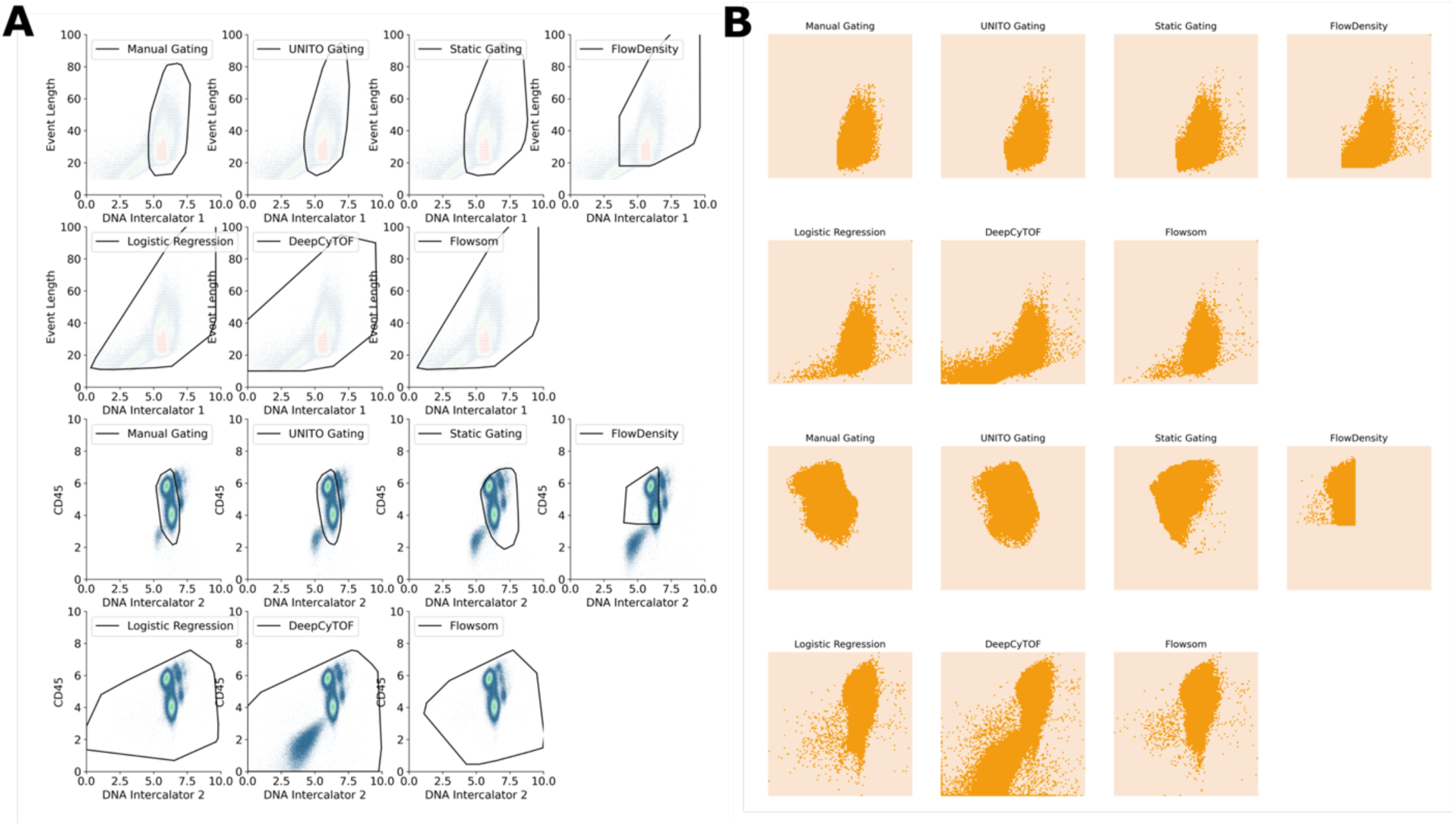
Gating results from different automated methods. **Panel (A)** shows the convex hull based on the cell-level cell type assignment, and **Panel (B)** shows the cells classified at a raw binary mask level on the density plot. For each panel, the top two rows present gating results for single cell gate 1, and the bottom two rows show the results for single cell gate 2.

### UNITO Gating on Flow Cytometry Data

The same visualization procedure was also repeated for the flow cytometry gating (**Figure 6A**; **Figure 3**). For automatic gating on flow cytometry, the UNITO framework still performs well in sequential gating tasks with both quantitative evaluation and visual inspection. The UNITO framework can accurately identify gates even when the biaxial map contains multiple, highly dense cell clusters, like in the side scatter-area (SSC-A) by foreword scatter-area (FSC-A) gate for lymphocytes (**Figure 6A, left**). Because of the presence of a much denser region of debris, the conventional semi-supervised algorithms always require some human-inspected hints into the pipeline to manually eliminate a certain region and give an approximate position of where the actual lymphocyte cluster may present. With limited data available, the UNITO framework can easily overcome this limitation and automatically eliminate the debris, disregard other clusters around, and only keep the lymphocytes cluster. The consistency of close-to-manual performance on subsequent gating proves UNITO’s ability to learn gating patterns for different cytometric data modalities. UNITO also maintains a good performance when comparing the proportion of target cells in the UNITO gating and manual gating (**Figure 6B**), in which all Pearson correlation coefficients are larger than 0.9.

**Figure 6.**
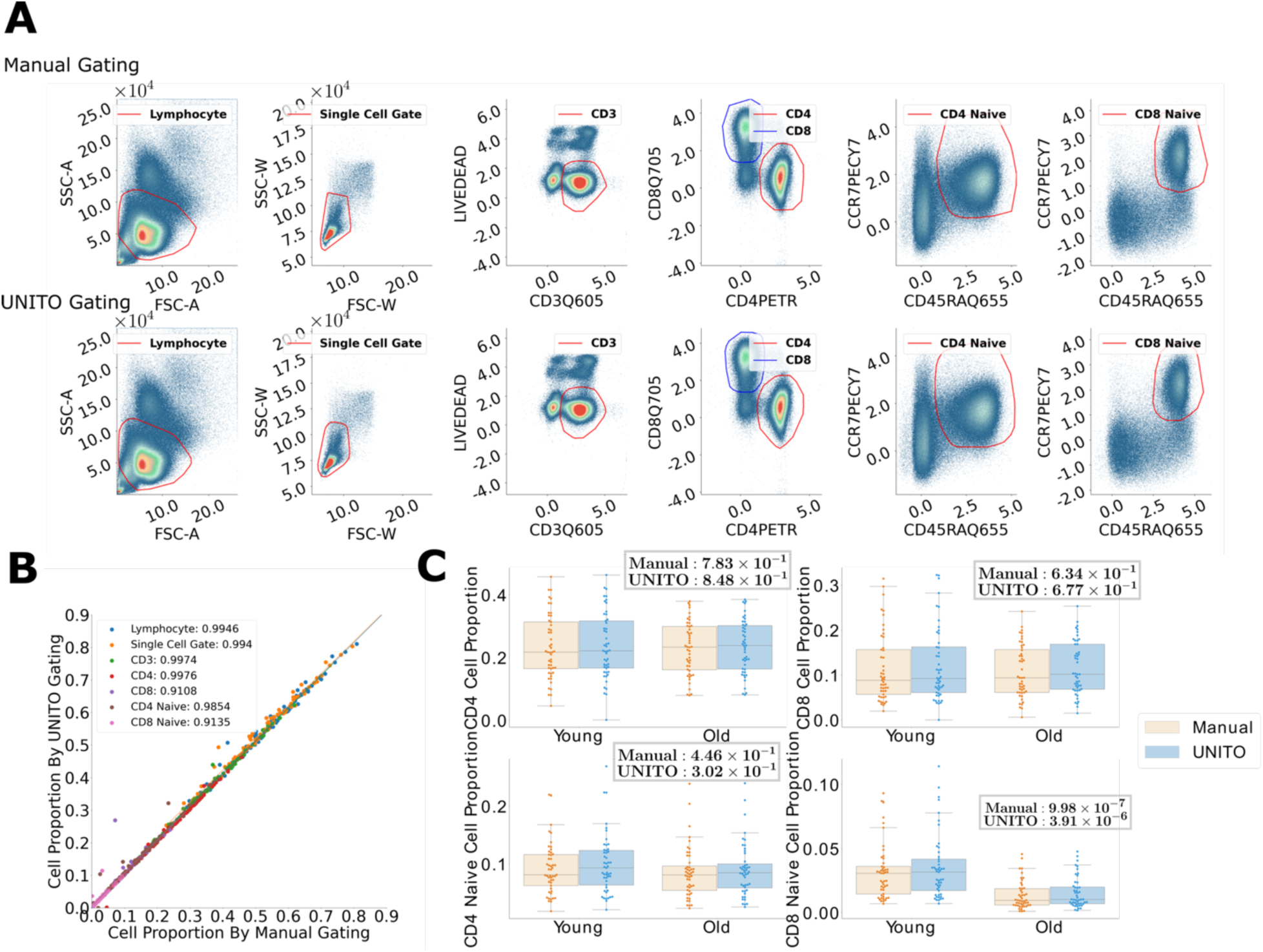
Comparison between UNITO automated gating and manual consensus gating (flow cytometry). **(A)** Density map and target cell annotation for flow cytometry data. The first row shows the ground truth annotation for one validation subject in the sequential prediction task, and the second row shows the predicted decision boundary from the reconstruction of the UNITO output. Each column represents a single level in the gating hierarchy, and gates within the same level are plotted together, such as CD4 cells and CD8 cells. **(B)** Visualization of the proportion of cells from UNITO and manual gating with all gating tasks together. **(C)** Statistical comparison validated on the Young vs Old flow cytometry data (Top row: CD4, CD8. Second row: CD4 naïve, CD8 naïve). The T-test between Young and Old group was applied on selected gates and P-values were reported for both manual gating and UNITO gating. For each gating, the plot is separated by subjects’ age condition while the results from manual gating and UNITO for the same population are put next to each other for easier observation.

### UNITO identifies immune health signatures by subtyping cell populations

Identifying similarities or differences among subjects in certain cell subpopulations is a common goal with the task of manual gating. To test whether the UNITO framework could support a workflow in which cell populations were compared between two groups, we analyzed their statistical difference between peripheral blood from healthy donors and patients with acute COVID-19 using the auto-gated cell data as well as the consensus-gated labels. An interesting property of UNITO is its ability to gate cells in a sequential manner, so the statistical tests can be applied either to the end nodes of the gating hierarchy tree or the middle nodes. Here we selected 6 gates after the pre-gating tasks to perform a t-test over the COVID population and healthy population (**Figure 4C**, **Figure S7**). We found that there are significantly different cell proportions between the two populations such as neutrophil, granulocyte, as well as lymphocyte, which agrees with previous literature^25–28^. The results show that the UNITO output after the pre-gating tasks can yield the same results among different cell sub-population tests compared to manual gating. In addition, the UNITO pipeline provided enough statistical power to distinguish group disparity between COVID and healthy subjects in many gates that are well-known to be drastically increased when facing immune diseases.

In addition, with the flow cytometry data, we also have the subject phenotype of young and old subjects (**Figure 6C**, **Figure S8**). While it is more difficult to observe the disparity among cell populations from young and old people than between COVID-19 patients and healthy donors, both UNITO (p-value 3.91e-6) and manual gating (p-value 9.98e-7) observe a significantly higher proportion of CD8 naïve cells in young donors as previously reported^29,30^. UNITO again provides enough statistical power as well as an equivalent gating p-value, even for the other cell types that were not significantly different between young and old donors.

## Methods

### UNITO Architecture

In this study, our UNITO framework was adapted from the architecture of the UNet model. UNet is a convolutional neural network (CNN) architecture that was first introduced by Ronneberger et al. in 2015^31^. The purpose of its original design was to perform semantic segmentation of medical images, which is the task of assigning a label to each pixel and thus defining the region of interest. The UNet architecture consists of two parts: an encoder (the contracting path) and a decoder (the expansive path). Both the pooling and up-sampling parts have a large number of feature channels, which allow the network to propagate context information to higher resolution layers which results in the expanding part being symmetric to the contracting path and yields a u-shaped architecture. The architecture of the encoder resembles that of a traditional CNN, which consists of a series of convolutional and pooling layers. The convolutional layers extract high-level features from the input image and the pooling layers reduce its spatial resolution. The decoder consists of a series of convolutional and transposed convolutional layers. The transposed convolutional layers upsample the feature maps to a higher resolution, which is achieved by interpolating the in-between pixels. The convolutional layers then process the upsampled feature maps to extract high-level features. At each decoding stage, the upsampled feature maps are concatenated with the corresponding feature maps directly passed from the encoding stage. These skip connections allow the decoder to use the contextual information from the encoder, which helps recover the lost spatial resolution.

We adapted the UNet architecture in our study to allow binary classification of each gate, and the output bivariate mask was used to produce the final segmentation image. Overall, the network consists of convolutional layers with hidden sizes from 1, 64, 128, 256, 512, and 1024, and then the transposed convolution will upsample the image back to the same size by following the same hidden size order but in the opposite direction. The skip connections provide the decoder with the original contextual information of the image, allowing U-Net to produce highly accurate segmentation results. In addition, U-Net has shown excellent performance in handling small data sets, which makes it particularly useful in medical applications, where the sample size is often limited.

### Cytometric Gating Structure

The main objective of UNITO is to achieve a similar boundary compared to human annotation with the ability to handle batch effects and heterogeneity between different subjects. Since subsequent cell-type gating tasks have the same setting as the pre-gating task, but with less variability, the original application of pre-gating cytometric data can be further extended to automatically gating cell subtypes in mass cytometry, as well as to gating flow cytometry data. **Figure S1A** shows all 5 levels of pre-gating and gating tasks (9 gates in total) of mass cytometry gates validated using UNITO. Each level from the hierarchy uses different pairs of channels from protein measurements, and each gate will undergo separate training. Within the same hierarchy, the cells are gated on the same space coordinate with the same pair of protein expressions, such as granulocyte and mononuclear. In addition to mass cytometry gating, **Figure S1B** shows the gating procedure for flow cytometry. Lymphocyte gate in flow cytometry data is still difficult to automate due to its high density of debris and noises from other cell populations. Thus, it usually requires manual gating or prior knowledge such as input of a hard threshold and approximate position based on the experiments. We validated the performance of UNITO starting from the lymphocyte gate and examined all the way to the naïve CD4+ and naïve CD8+ T cells.

### Construction of Bivariate Density Map and Mask

Here we define our cytometric data *X_c_* = {*X*_1_, *X*_2_, *X*_3_, …, *X_n_*} where each *X_i_* represent a single subject. Let *Xi* = [*x_uv_*] ∈ ℝ*^m*n^* be the protein expression matrix, where *m* is the number of cells in the experiments and *n* is the number of channels collected. Usually, a specific cell distribution mostly varies among two protein measurements, and using two channels for manual gating is easier to visualize and inspect. Therefore, two selected cytometric measurements are normalized and rounded as a constraint on the boundaries for the density map and mask. The normalization process is defined that for each *x_uv_* in the cytometric measurement *X*:

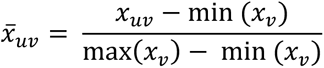

Density maps of the normalized cytometric data were created as training images by counting the number of cells with the same pair of normalized measurements and feeding the number into the corresponding coordinate in the matrix space. In this case, the density matrix has corresponding shape of 101 by 101 representing the normalized measurement. This transformation not only made the process of creating density maps easier, but also allowed faster training and prediction time because this step reduced the training input from a data matrix of millions of entries to a 101 by 101 matrix. The training data also provides gating labels for each cell inferred from manual annotation, and the corresponding binary masks were created from the manual labels as the training labels. If the cells are identified as the target population, the corresponding coordinate will be marked as 1, creating a 101 by 101 binary image. In addition, a convex hull algorithm was further applied on the binary mask to smooth the boundaries using the ConvexHull function from the Python package SciPy^32^.

### UNITO Training and Cell Membership Prediction

The UNITO was trained using the PyTorch framework in Python. We searched hyperparameters learning rate and batch size for each gate, and we also checked convergence during each training process. The Adaptive Moment Estimation Optimizer^33^ or in short, ADAM optimizer is a popular optimization method that allows the learning rate to adjust to the gradient change during the training process adaptively. This along with the popular Binary Cross-Entropy Loss function with Logits (BCEWithLogitsLoss) is used in the binary classification of the pixels. The Binary cross-entropy Loss can provide a good gradient calculation for optimization by combining the sigmoid function and binary cross-entropy loss (BCELoss) into a single function and is more numerically stable than using a plain sigmoid followed by a BCELoss.

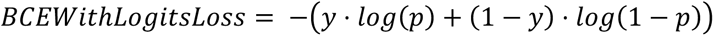

Once we have the trained model, we can use the pipeline to predict the binary labels for each cell. Given a target subject, we first use its normalized cytometric data to create its density map corresponding to its gate. This density map also has a resolution of 101 by 101 and is obtained using the above-described procedure. It is then passed into the trained model to generate the predicted mask for the assigned gate. This binary mask is then interpolated with the protein measurement to generate the binary label for each cell. Specifically, each cell corresponds to a pixel on the binary mask; if a pixel is predicted as in-gate, all the cells corresponding to the selected measurements will be designated as in-gate, and vice versa. Next, we filter out all the cells that are not predicted to be in this gate, and we use the remaining data (in-gate cells for the current gate) to create the density map corresponding to the next gate.

Again, the pre-processing and training procedure will be repeated for the next gate using the filtered cytometric data. While all gates are trained and all subjects are predicted, we evaluated the performance of each gate and visually inspected the bivariate polygons on the density map. We can also use intermediate or final gate results to perform downstream statistical analysis. The details of the UNITO algorithm are summarized in **Algorithm 1**.

**Algorithm 1:**
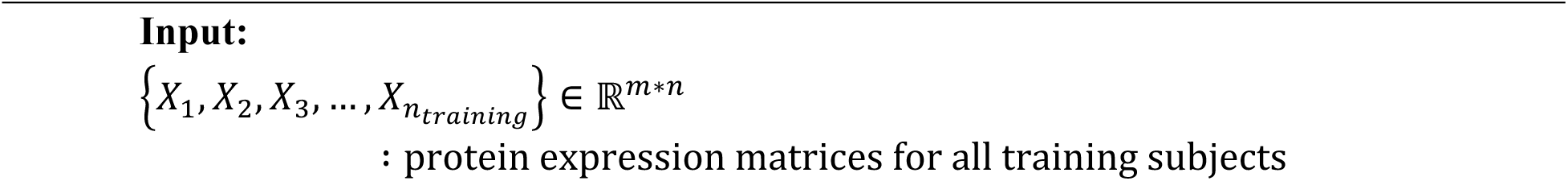
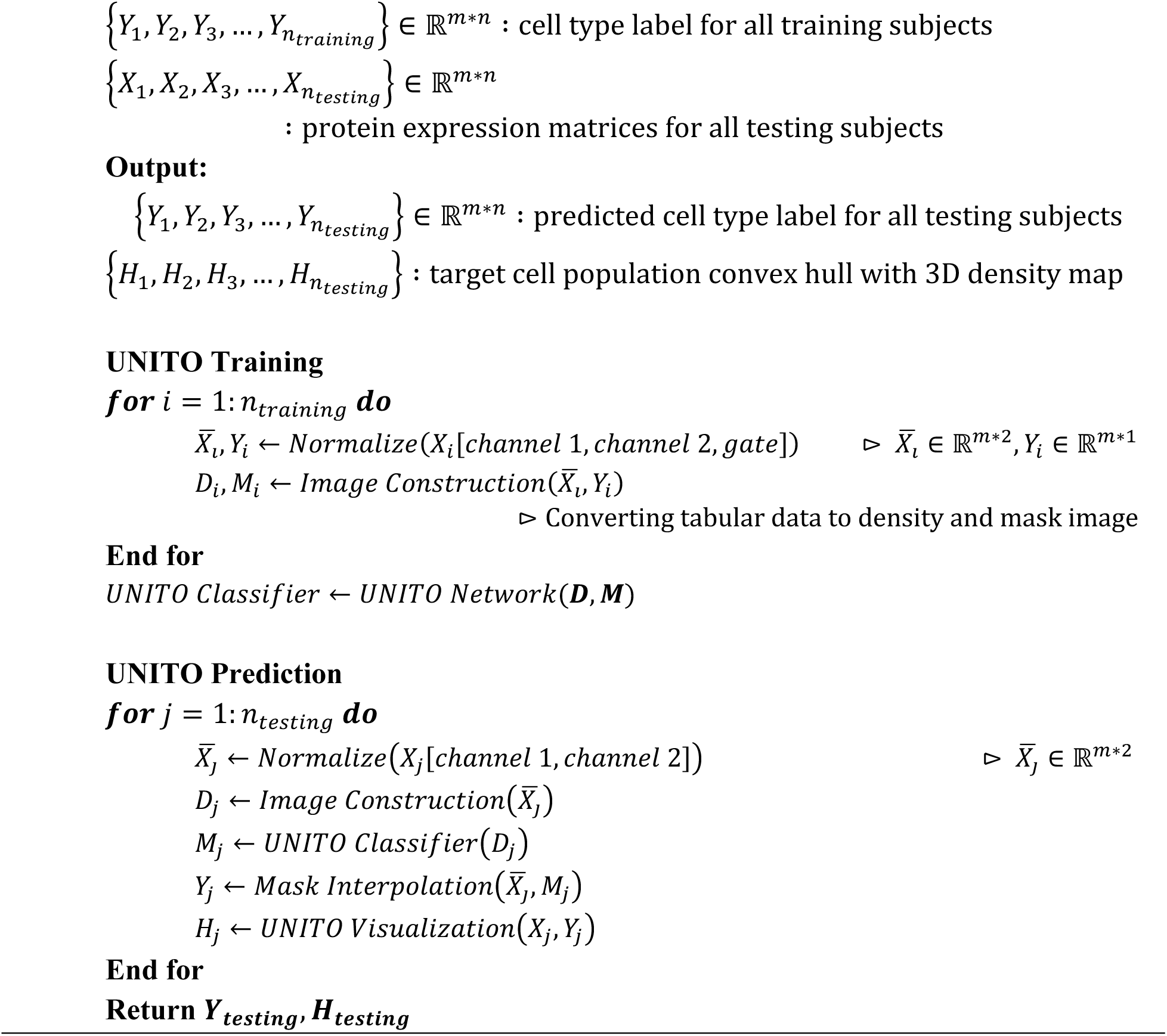
UNITO Implementation Input:

### Mass Cytometry Data Collection and Processing

We used two independent datasets consisting of single-cell protein expression data profiled by mass cytometry (**Table 3**). Both datasets were collected by the Institute for Immunology and Immune Health at the Perelman School of Medicine. For the first (“COVID Vaccine”) dataset, whole blood was obtained from 40 healthy subjects at four timepoints during the course of two-dose mRNA vaccination against COVID-19 and cryopreserved. (T1 = baseline, T2 = one week after the first dose, T3 = prior to the second dose, T4 = one week after the second dose). Only the baseline sample was used, in order to avoid information leaks across training. For the second (“COVID-19/healthy”) dataset, whole blood was obtained from 23 subjects, among whom 12 had acute COVID-19 symptoms and 11 were healthy donors. Samples collected for the COVID-19/healthy dataset were used fresh.

**Table 3.** Demographic details of three datasets. The table shows the number of subjects and cells for each small group in three datasets. The total number of cells is reported in the format of (mean ± standard deviation).

|  | Mass Cytometry Dataset |  |  | Flow Cytometry Dataset |  |
| --- | --- | --- | --- | --- | --- |
|  | COVID Vaccine Dataset | COVID vs Healthy Dataset |  | Young vs Old Dataset |  |
|  |  | COVID subjects | Healthy subjects | Young Subjects | Old Subjects |
| Subjects | 40 | 12 | 11 | 49 | 48 |
| Cells | 321k ± 156k | 304k ± 228k | 339k ± 215k | 1092k ± 372k | 953k ± 291k |

Both the fresh samples from the Acute dataset and the frozen samples from the Vaccine dataset were stained with the Maxpar Direct Immunophenotyping Assay (MDIPA) and run on a CyTOF2 instrument. Samples were stained, stored, and prepared for acquisition as previously described ^34^. MDIPA measures a panel of 30 proteins used for broad characterization of immune phenotypes in whole blood, alongside other channels for control and data cleaning. Raw CyTOF data was transformed using an asinh transformation with a cofactor of 5. Further, a standard data cleaning procedure by manual gating was performed using the OMIQ platform, to remove beads, dead cells, platelets, debris, and other anomalous events, followed by manual gating on the subsequent cell type to generate the cell type labels for this study.

### Flow Cytometry Data Collection and Processing

The young/old dataset was created by Rochester Human Immunology Center, David H. Smith Center for Vaccine Biology and Immunology, Rochester, NY (USA)^35^. The purpose was to use SWIFT’s competitive clustering assignment method to measure the differences between PBMC sub-populations in Old/Young subjects. The Young/Old dataset contains 136 data files in total, with 97 unique subjects; similarly to the COVID Vaccine dataset, only one file from each subject was used, to avoid information leaks. Within those subjects, there are 49 samples from young donors and 48 samples from old donors. The dataset was downloaded from the publicly available repository (https://flowrepository.org/id/FR-FCM-ZZGS). The data cleaning and labeling by manual gating were also done using the OMIQ platform.

### Manual Gating and Consensus

The mass cytometry and flow cytometry data were manually gated by 5 and 4 independent annotators, respectively (see **Figure 4** for mass cytometry data and **Figure 5** for flow cytometry data). The final version of the label used for training and validation was generated by the consensus (majority vote) of the gating output from all annotators. Specifically, for each cell, we calculated the highest frequency of the cell type assignment from the different labels and saved the results as our final consensus voting label. **Figure 1** and **Figure 6** shows the correlation between consensus voting labels with manual gating from each annotator.

## Discussion

The rapid development of single-cell technologies has enabled large-scale and high throughput data collection, but at the same time raised challenges for analyzing such large amounts of data. The size of data not only increased within a single sample (number of cells), but we are also accumulating more donors over time. In this study, we first described a common challenge in cytometric pre-gating tasks due to the high technical and biological variability across subjects. Most state of the art unsupervised auto-gating methods typically assume the data is already cleaned from the pre-gating stage and do not explicitly handle events such as debris and doublets, which are unpredictable and difficult to model. Other auto-gating methods such as FlowDensity also require human prior knowledge as hints for the model to target the position or percentage of the desired cell subpopulation. There are also supervised methods that predict cell types using deep neural networks. However, treating every single cell as one data point will require large computational resources to both train the model and predict incoming data. The proposed framework UNITO transformed the numerical classification of protein expression data into an image-based semantic segmentation task. In this case, by using the convolutional kernel to target the densest region in the cell density plot, the UNITO framework guaranteed the translational invariance property of cell clusters and a global view of the bivariate cell density distribution. With such properties, the proposed method can be applied and extended to any cytometric data including mass cytometry and flow cytometry to automatically gate for multiple purposes, such as removing debris, doublets, and gating other cell subtypes. By converting the protein expression matrix to the density map, UNITO further allows the incorporation of more training data and is capable of learning the protein expression behavior in a systematic pattern across subjects. One advantage of UNITO is that since the prediction is performed in each single sample file by file, it can query a huge number of samples parallelly with very low computational cost, even computing with CPU is sufficient for prediction. In addition, the computing speed of UNITO will also not be affected by the number of cells in each sample, which guarantees its efficacy even if the throughput of the cytometric experiments continues to grow in the future. Using UNITO for immunophenotyping in different protein panels would require separate training. However, cleanup gates such as Single cell gate 1 and 2 in CyTOF, and the Lymphocyte and Singlet gates in flow cytometry, are the same in most experiments. Therefore, researchers can use the models we already trained to perform cleaning on their own data.

The UNITO performance on the sequential pre-gating and gating tasks for immune cells proved the similar-to-human boundaries of the UNITO framework by visually comparing the boundaries of human annotation and UNITO prediction on the density map. One of the biggest advantages of the UNITO framework is that it also gives the boundary of the target cell population, which enables the interpretability of the gating results and allows inspection of intermediate gating steps. While the cell-level prediction using logistic regression or deep neural networks and unsupervised clustering also gives good evaluation scores, the reliability of the gating results with biological knowledge is still in debate. We visualized the gating results with an example subject for the first two single-cell gates in the mass cytometry data (**Figure 5**), and we can clearly see that only UNITO has the closest contour compared to the manual gating. The static gating has a similar shape, but the position is off due to the biological variability of the subject and the gating position is fixed. The idea of FlowDensity is to segment the entire cell population into four quadrants in the coordinate space and based on the position parameter and percentage to draw the boundaries. In this case, we can observe a clear right angle on the contour. In our FlowDensity group-prediction experiments, we tuned and fixed those parameters for each gate, so the area of boundaries might be too large or too small for certain subjects. This is due to the fact that in reality the percentage can vary a lot across subjects (**Figure 2,3,7,8**). Another alternative called flowLearn^21^ uses an algorithm instead of a hyper-parameter to find the best cutoff and perform the density-based gating, but the implementation of only relying on one protein measurement at a time to separate two peaks makes automatic pre-gating difficult. The visualization of gating results on the density map for logistic regression and DeepCyTOF almost included the entire coordinate space, indicating that cell-level predictions based on high dimensional protein measurements are difficult to validate in the bivariate setting familiar to immunologists. Even though those methods have high evaluation scores, they will always include cells that are far away from the desired cell population. A similar situation also happens in the FlowSOM clustering, where the convex hull included a large region with a noisy pattern in the binary mask visualization (**Figure 5**). A simple observation we can draw from this comparison is that prediction over bivariate image outperforms prediction on single-cell tabular data in both accuracy and interpretability, and even static gating showed its capacity to provide a good gating prediction. If there are large batch effects in the data, the performance of static gating will decrease, but UNITO is still robust to such variability.

The key goal for all of the pre-gating and gating tasks is to discover the hidden information embedded in the cell subpopulations. The downstream analysis usually reveals statistical differences either among cell subtypes or subject groups. With prior knowledge of the biological difference between COVID-19 patients and healthy donors, as well as between young and old people, we can further use the statistical power of the gating results to validate the method’s performance. The proportion of cell types selected by prior knowledge in UNITO gating confirmed a significant increase in granulocytes and neutrophils with a significant decrease in lymphocytes, T cells, and B cells, therefore validating the efficacy of UNITO. The same experiment on the flow cytometry data in CD8 naïve T cells also agreed with biological prior knowledge. This downstream analysis is not only used for confirming the gating results from UNITO, but also serves as a functionality to explore the group-level difference for scientific discovery purposes.

There are certain limitations to this study. The model may not get very good performance in certain cases where cells are extremely rare, such as eosinophils in certain samples. Even though the UNITO framework still has the highest performance for eosinophil prediction, we found one file in which there were almost no eosinophils. In this case, even in the manual gating, eosinophils can barely be seen. For such situation, one mitigation is to not only pre-filter the cells by the previous gating level, but also pre-filter the other cell types in the current gating level. For instance, after we filtered out the non-granulocyte cells, we can add additional steps to filter predicted neutrophil cells, leaving only eosinophil and non-neutrophil-non-eosinophil cells in the data to segment the eosinophils. Moreover, we only tuned hyperparameters for the learning rate and batch size when designing UNITO, more advanced deep learning techniques such as adding self-attention blocks or using different image segmentation backbones may improve the performance.

## Conclusion

To summarize, we present a new framework that can automatically perform pre-gating and gating tasks for both mass cytometry and flow cytometry with close-to-manual performance. We validated that autogating by bivariate images outperforms gating on cell-level protein expression data, and UNITO can further provide the convex boundary on the density map for h1biological interpretation. In addition, UNITO predictions are easy to use in downstream statistical analysis for cell type-phenotype exploration, and yield results that are as statistically significant as those from manual gating. We then showed that the framework can be adapted to any subsequent gating tasks in a sequential manner while still maintaining high performance. Future work can extend this framework to develop a fully automated end-to-end data processing pipeline for cytometric data. Originally developed in Python, leveraging its seamless integration with deep learning packages like PyTorch, UNITO currently requires users to run scripts with specific gating parameters via terminal or scripts, using pre-processed CSV files derived from FCS files typical in cytometric experiments. To enhance the accessibility and user-friendliness of UNITO, our future work includes the development of a web-based or software-based iteration taking raw FCS files as input, which is in progress with the Pennsieve Data Management Platform from the University of Pennsylvania.

## Author Contributions Statement

J.C. conceptualized and developed the model, wrote the source code, and wrote the paper. N.M coordinated the COVID-19 study, recruited subjects and collected clinical data. M.M, M. P, D.M, A.P, A.G collected and processed the data. J.C., M.I., Y.F., and Y.L. conducted relevant computational experiments. M.M and M.I developed the framework for aggregating and comparing results from multiple annotators. A.P, M.P, M.M, D.M, M.I manually annotated data. All authors reviewed the paper. L.S. supervised the study and reviewed the method.

## Acknowledgments

This work was supported in part by the following NIH grants: U01 AG068057, S10 OD023495, and the Glick COVID-19 research award. We thank Takuya Ohtani and the CyTOF Core at the University of Pennsylvania for data acquisition.

## Supplementary Material

### Supplementary Notes

#### Comparison Methods

We used k-fold cross-validation to ensure that all samples from each dataset get their prediction iteratively. For the mass cytometry data, we have 63 subjects in total, and we divided the cohort into 8 subsets (7 of them having 8 subjects and the last one having 7 subjects). For each iteration, we train on 7 of the 8 subsets and predict on the remaining set. Similarly, for the flow cytometry data with 97 subjects, we divided the dataset into 10 folds, with 10 subjects in each fold and 7 subjects in the last fold. UNITO and all comparison methods were trained and predicted on the same subset of subjects to ensure the consistency of the comparison.

#### Static Gating

Static gating creates a consensus mask from all training subjects and uses that as a fixed mask for all testing subjects. During the training process, static gating uses the same pre-processing as UNITO, where the selected protein measurements are normalized as follows:

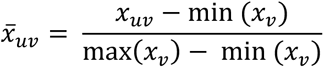

The bivariate masks for each subject in the training subset are created based on the normalized expression measurement. Then the average over all training masks was calculated to get a consensus mask, and a threshold was applied to binarize the bivariate mask. We tuned the threshold from 0.1, 0.2 to 0.9 for all gates separately to find the optimal static gates. Density maps were created only for testing subjects, and the thresholded bivariate mask was then applied to each testing subject to evaluate the performance.

#### FlowDensity

We used the R package FlowDensity (v1.35.0)^1^ from Bioconductor. There are two parameters we need to decide on, which are the position parameter and the percentage parameter. Each parameter is a tuple containing two elements corresponding to two dimensions in the input data respectively. For the position parameter, we only have True or False representing the relative position in the density-segmented space, and the percentage controls how large the region of interests should be in the density plot. Since we have multiple gates and multiple folds to train, it is very time-consuming to tune all combinations over the position and percentage parameters. Therefore, we first fixed the percentage to 0.5 for both axes and tuned all four possible pairs of position parameters, and after we got the optimal position parameter, we then tuned the percentage parameter, which ranges from 0.1, 0.2 to 0.9, in total results in 81 possible pairs. Finally, we use the obtained optimal parameter to predict the region of interest for the remaining testing subjects and convert that back to the cell-level labeling.

#### FlowSOM

We used the R package FlowSOM (v2.6.0)^2^ from Bioconductor. Data from all subjects and all channels necessary for the gating scheme under consideration (10 channels for CyTOF, 9 channels for flow cytometry) was pooled together into one matrix and analyzed with FlowSOM. For CyTOF, the two DNA intercalator channels were first scaled by the value of the 95^th^ percentile in each file, as a heuristic to account for the large inter-sample variability of these channels. We used the default settings of a 10 by 10 grid and 10 final clusters for CyTOF. For flow cytometry, we increased the number of final clusters to 30, because smaller numbers did not produce a good enough resolution to enumerate all cell types of interest. Each of the final clusters was labeled a belonging to a gate in the hierarchy based on the location of its centroid (mean of all events contained in the cluster). We remark that FlowSOM is an unsupervised method and makes no use of the labeled training data. On the other hand, its performance is very sensitive to the choice of data normalization and final number of clusters, as described above.

#### Logistic Regression

For the cell-wise classification task, we use all features selected in the entire gating structure as input to predict the cell type. We keep the idea of the hierarchical order but do the prediction in one step. Specifically, there are 8 classes in the prediction, which are eosinophil, neutrophil, t cell, b cell, non t non b cell, granulocyte (apart from all eosinophils and neutrophils), mononuclear (apart from all T cells, B cells and non-T non-B cells), single cell 2 (apart from all granulocytes and mononuclear cells), single cell 1 (apart from single-cell-2), and non-single-cell-1 cell. After we do the prediction, we integrate the cell type from the bottom up to higher level cells. For example, the final label of granulocyte gets all cells from cells with label granulocyte, eosinophil, and neutrophil. We used the LogisticRegression function from the Python Scikit-Learn package, and we tuned hyperparameter of the regularization with l1 and l2 norm, and with coefficient among np.logspace(−1,1,5), which is [0.1, 0.316, 1, 3.16, 10]. The prediction was done at all cells from the testing subset, and then we divide the cell by subjects for evaluation.

#### DeepCyTOF

The DeepCyTOF^3^ uses the same setup as the logistic regression, but with a deep learning method. We modified the code from https://github.com/KlugerLab/deepcytof.git to fit our data. Firstly, it only takes one file for training the cell classifier, so we integrated all files from training folds, which resulted in a large CSV file near 8 GB. In this case, normal GPU cannot handle such large dataset along with multiple large models due to limited GPU memory. In this case, we ran the analysis on a high performance cluster with an A100 GPU (80 GB of GPU memory). Other than that, we used the exact same neural network architecture and hyperparameter for the DeepCyTOF.

In addition, we also tested the DGCyTOF^4^ code from https://github.com/lijcheng12/DGCyTOF/tree/main. With the same setup and configuration, we moved the same neural network to our data to do the training and prediction. Unfortunately, the network did not converge so we did not report the performance in our manuscript.

### Supplementary Table

**Table S1.**
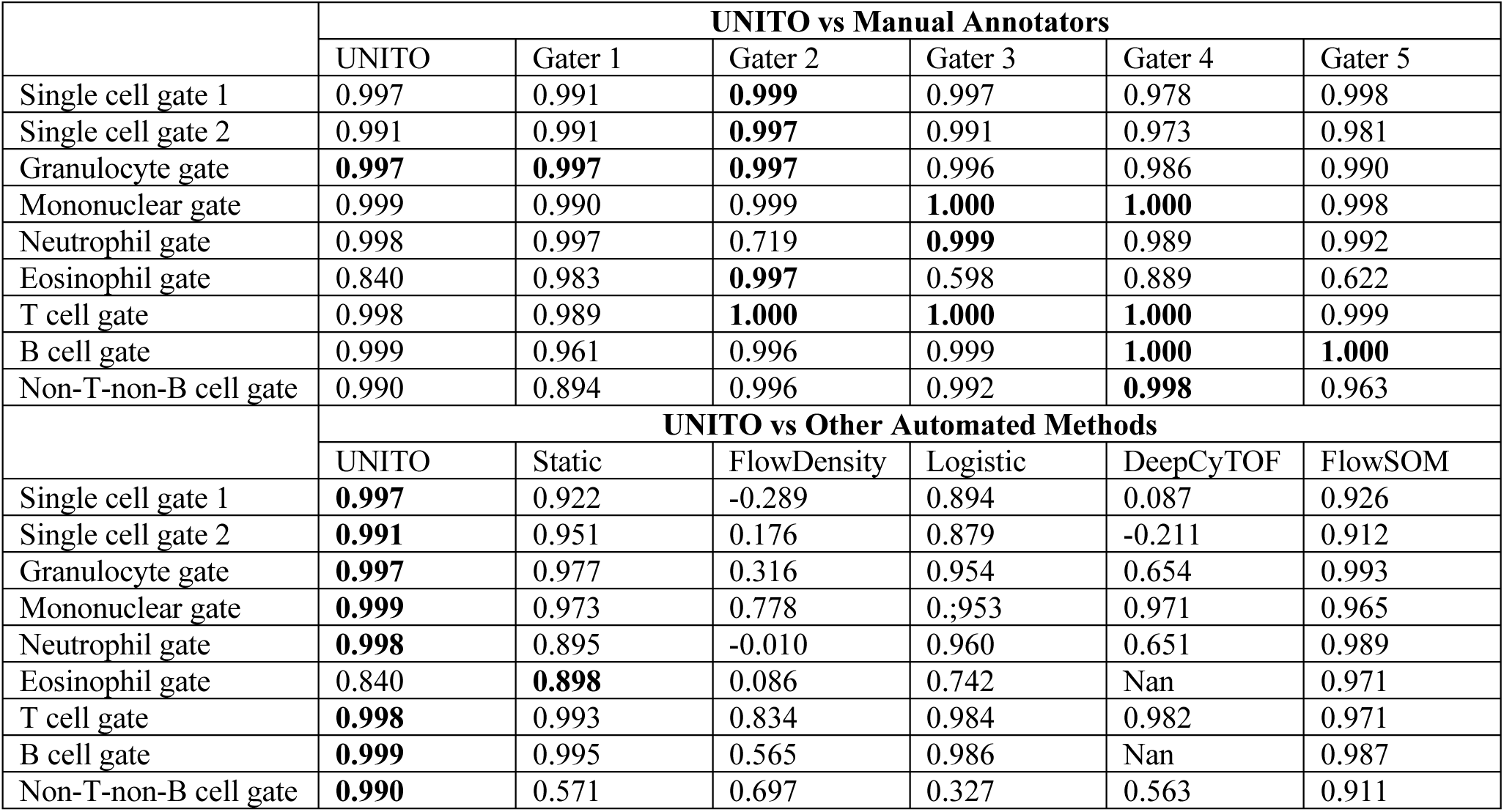
Pearson Correlation between percentage of gated cells from manual gating (consensus) with human annotator / automated methods in mass cytometry experiments.

|  | UNITO vs Manual Annotators |  |  |  |  |  |
| --- | --- | --- | --- | --- | --- | --- |
|  | UNITO | Gater 1 | Gater 2 | Gater 3 | Gater 4 | Gater 5 |
| Single cell gate 1 | 0.997 | 0.991 | **0.999** | 0.997 | 0.978 | 0.998 |
| Single cell gate 2 | 0.991 | 0.991 | **0.997** | 0.991 | 0.973 | 0.981 |
| Granulocyte gate | **0.997** | **0.997** | **0.997** | 0.996 | 0.986 | 0.990 |
| Mononuclear gate | 0.999 | 0.990 | 0.999 | **1.000** | **1.000** | 0.998 |
| Neutrophil gate | 0.998 | 0.997 | 0.719 | **0.999** | 0.989 | 0.992 |
| Eosinophil gate | 0.840 | 0.983 | **0.997** | 0.598 | 0.889 | 0.622 |
| T cell gate | 0.998 | 0.989 | **1.000** | **1.000** | **1.000** | 0.999 |
| B cell gate | 0.999 | 0.961 | 0.996 | 0.999 | **1.000** | **1.000** |
| Non-T-non-B cell gate | 0.990 | 0.894 | 0.996 | 0.992 | **0.998** | 0.963 |
|  | UNITO vs Other Automated Methods |  |  |  |  |  |
|  | UNITO | Static | FlowDensity | Logistic | DeepCyTOF | FlowSOM |
| Single cell gate 1 | **0.997** | 0.922 | -0.289 | 0.894 | 0.087 | 0.926 |
| Single cell gate 2 | **0.991** | 0.951 | 0.176 | 0.879 | -0.211 | 0.912 |
| Granulocyte gate | **0.997** | 0.977 | 0.316 | 0.954 | 0.654 | 0.993 |
| Mononuclear gate | **0.999** | 0.973 | 0.778 | 0.953 | 0.971 | 0.965 |
| Neutrophil gate | **0.998** | 0.895 | -0.010 | 0.960 | 0.651 | 0.989 |
| Eosinophil gate | 0.840 | **0.898** | 0.086 | 0.742 | Nan | 0.971 |
| T cell gate | **0.998** | 0.993 | 0.834 | 0.984 | 0.982 | 0.971 |
| B cell gate | **0.999** | 0.995 | 0.565 | 0.986 | Nan | 0.987 |
| Non-T-non-B cell gate | **0.990** | 0.571 | 0.697 | 0.327 | 0.563 | 0.911 |

**Table S2.**
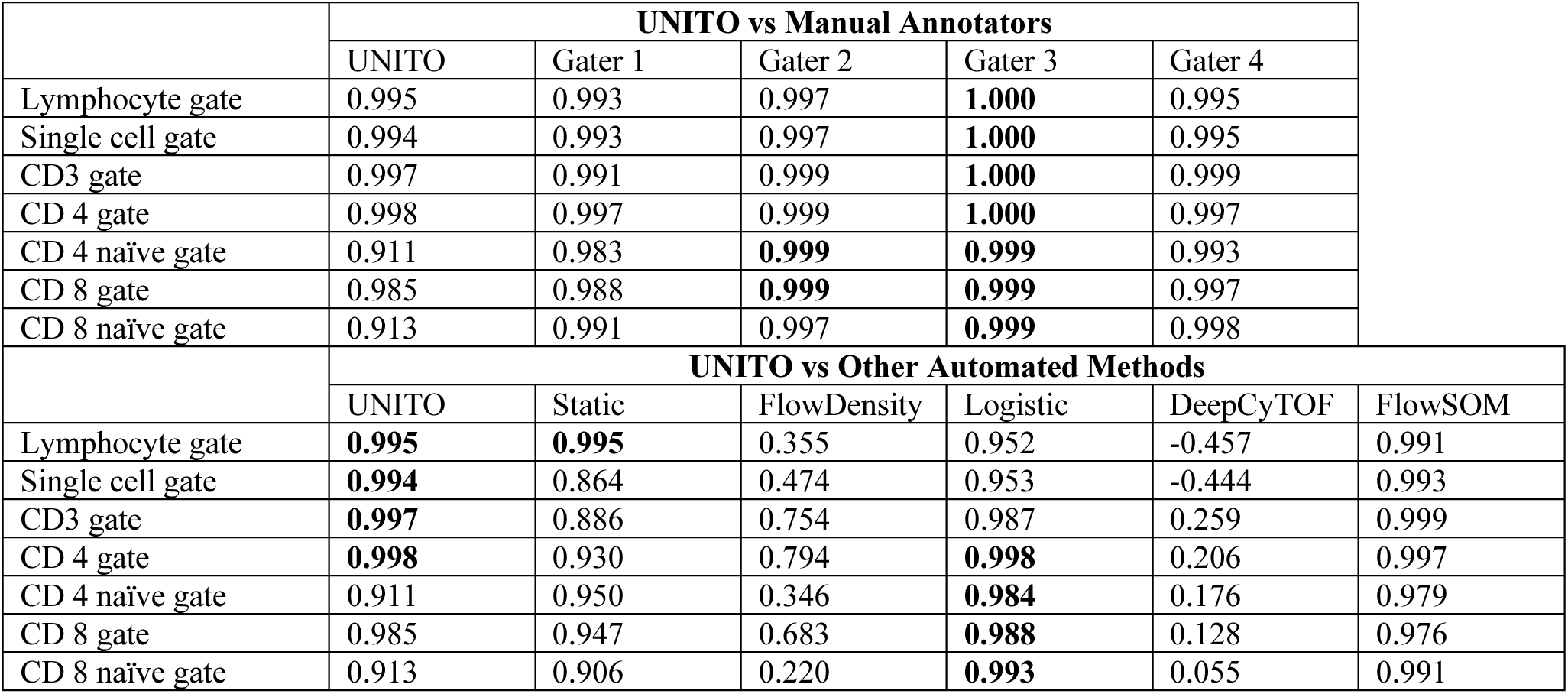
Pearson Correlation between percentage of gated cells from manual gating (consensus) with human annotator / automated methods in flow cytometry experiments.

### Supplementary Figures

**Figure S1.**
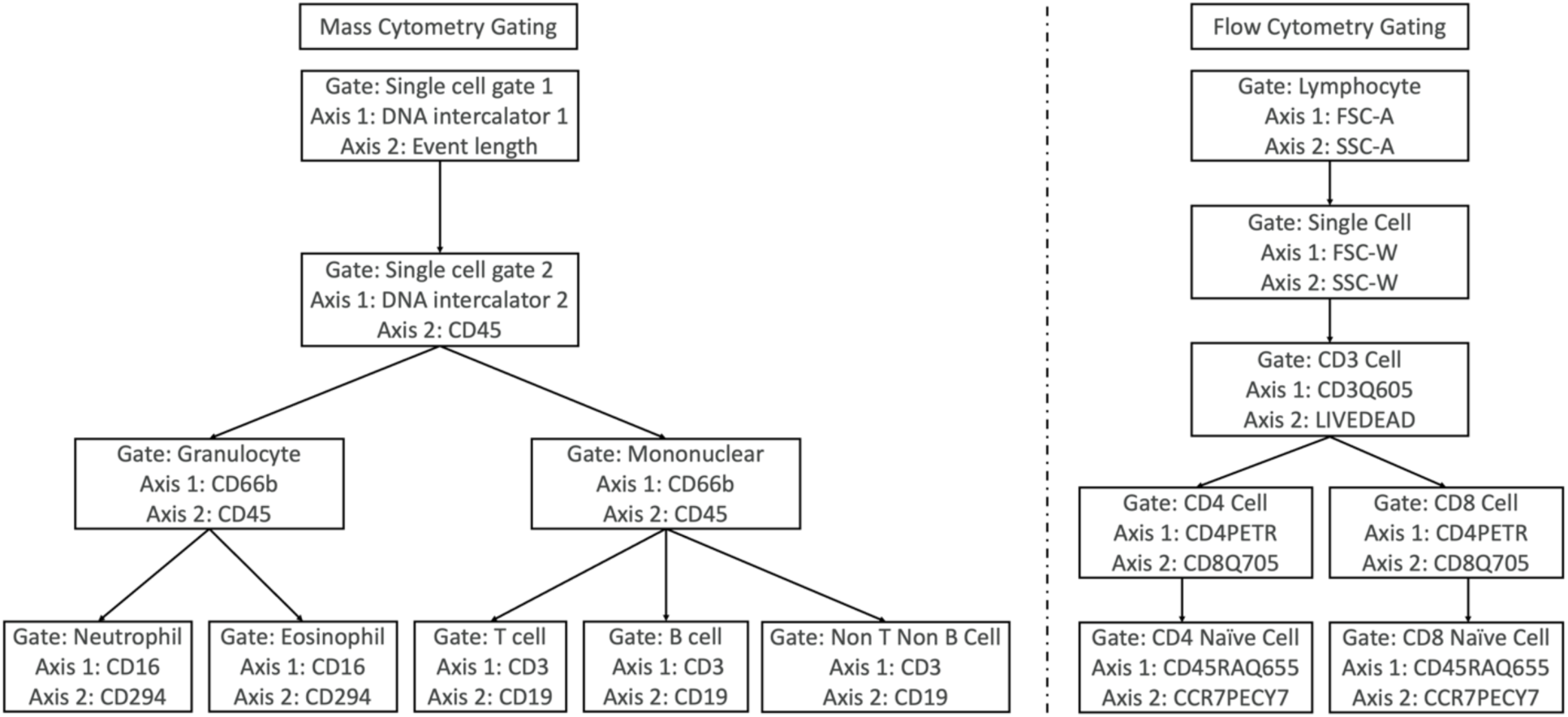
Hierarchical illustration of cytometric data pre-gating and gating structure. **A)** Mass cytometry datasets have four levels in gating tasks with 9 gates in total. While the pre-gating 1 and pre-gating 2 are sequential, subsequent gating are expended in a tree structure. **B)** The flow cytometry gating has 5 levels with 7 gates in total.

**Figure S2.**
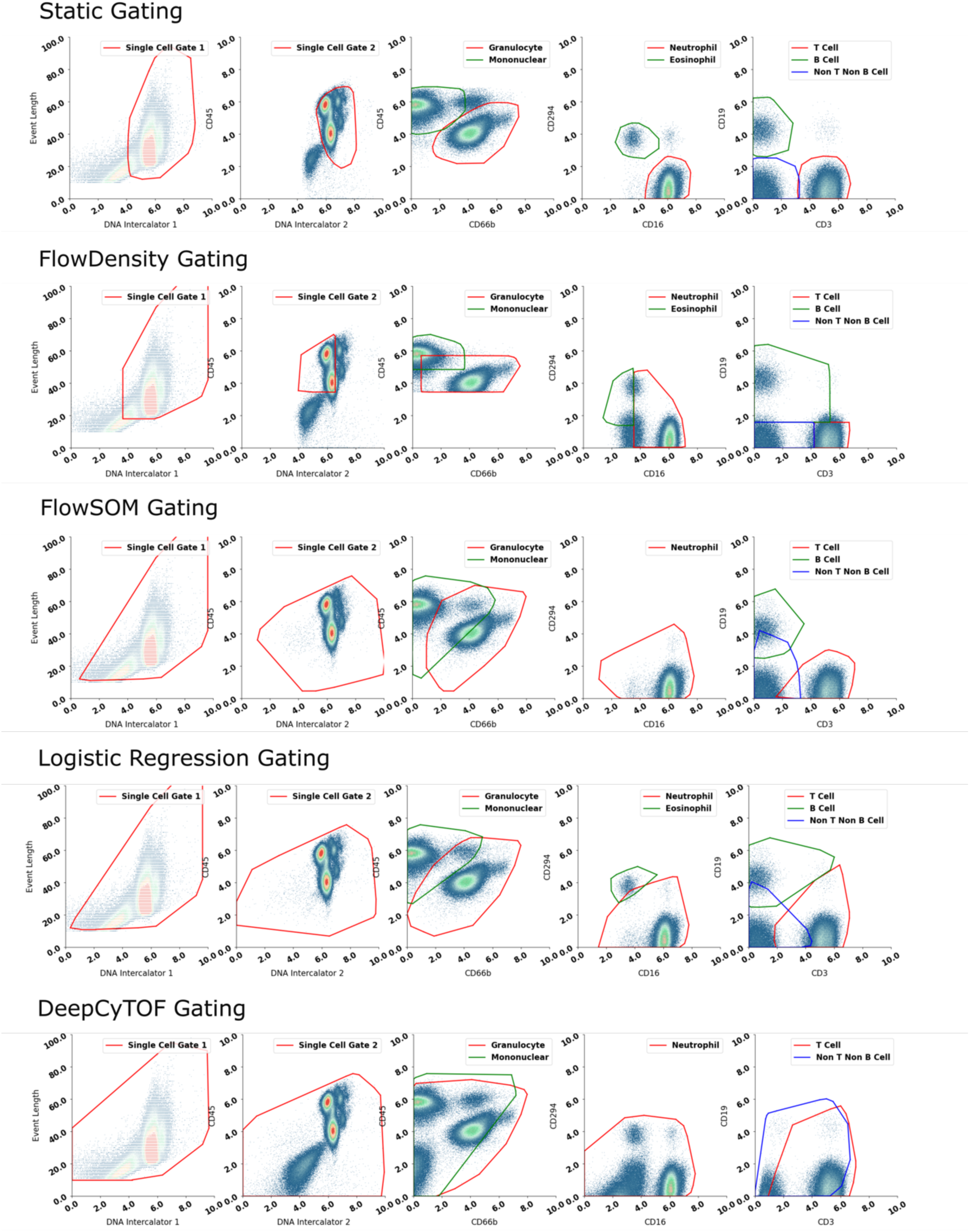
Gating visualization for all comparison methods (mass cytometry). Each row represents a comparison method (top to bottom: static gating, FlowDensity, FlowSOM, logistic regression, DeepCyTOF). Gates using the same protein channel are plotted in the same subplot (left to right: single cell gate 1, single cell gate 2, granulocyte + mononuclear, neutrophil + eosinophil, t cell + b cell + non t non b cell).

**Figure S3.**
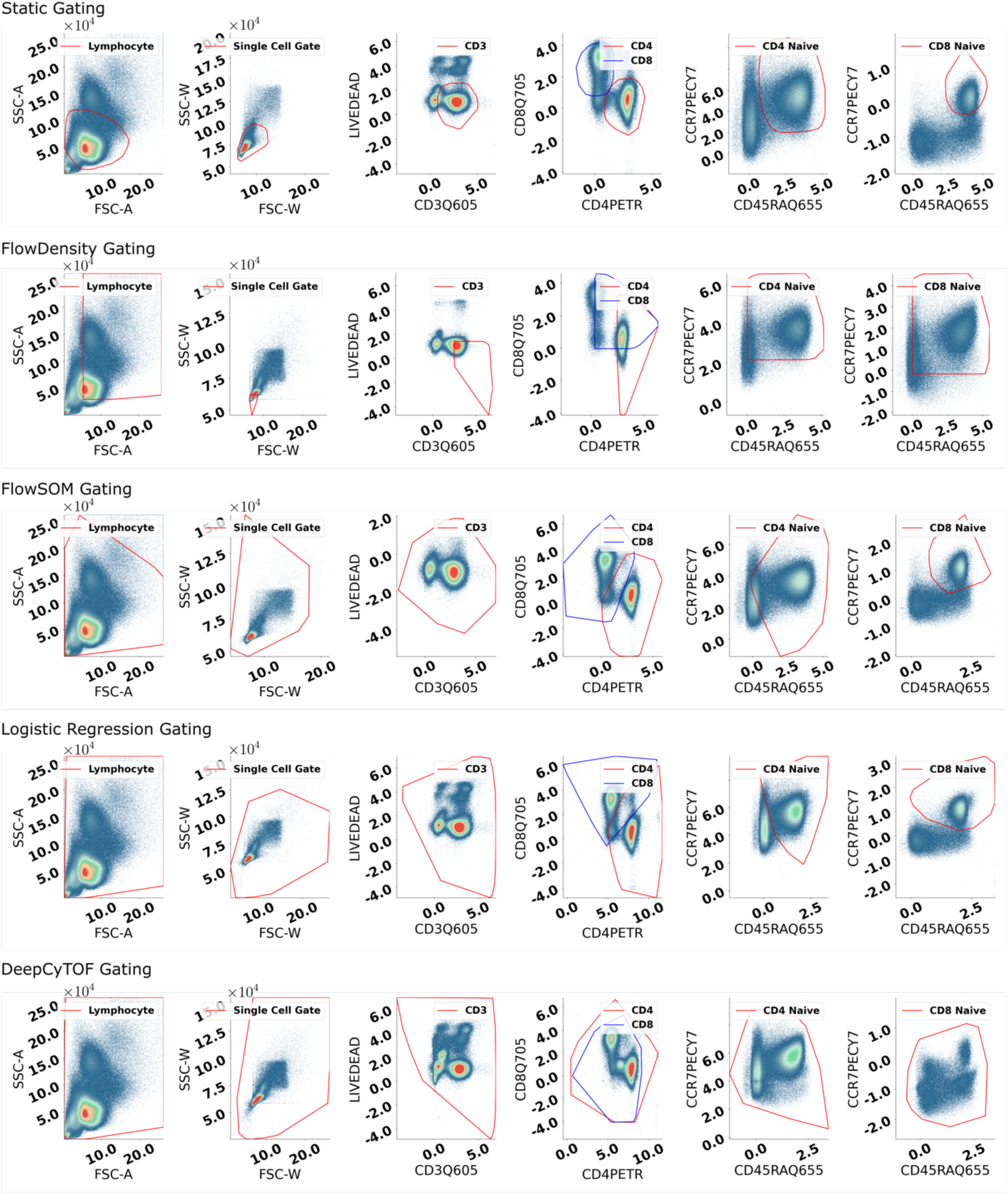
Gating visualization for all comparison methods (flow cytometry). Each row represents a comparison method (top to bottom: static gating, FlowDensity, FlowSOM, logistic regression, DeepCyTOF). Gates using the same protein channel are plotted in the same subplot (left to right: lymphocyte, single cell gate, CD3, CD4 + CD8, CD4naive, CD8naive).

**Figure S4.**
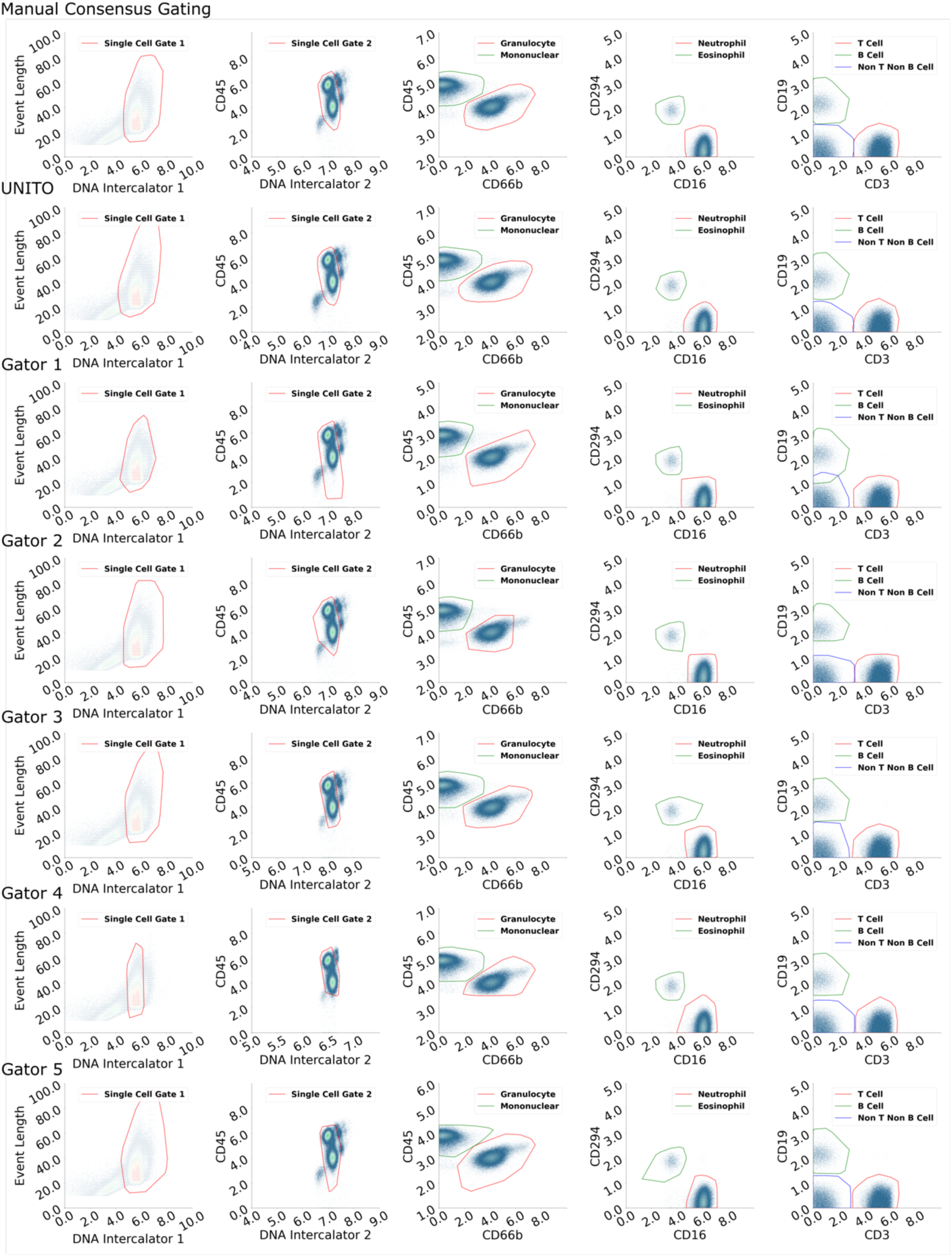
Gating results by gaters (mass cytometry). To assess the performance of UNITO compared to each manual gater, we plotted the consensus manual gating, UNITO gating, along with all other individual manual gating together (top to bottom: consensus manual gating, UNITO gating, gater 1-5). The gates are organized in the same way as previous figures (left to right: single cell gate 1, single cell gate 2, granulocyte + mononuclear, neutrophil + eosinophil, t cell + b cell + non t non b cell).

**Figure S5.**
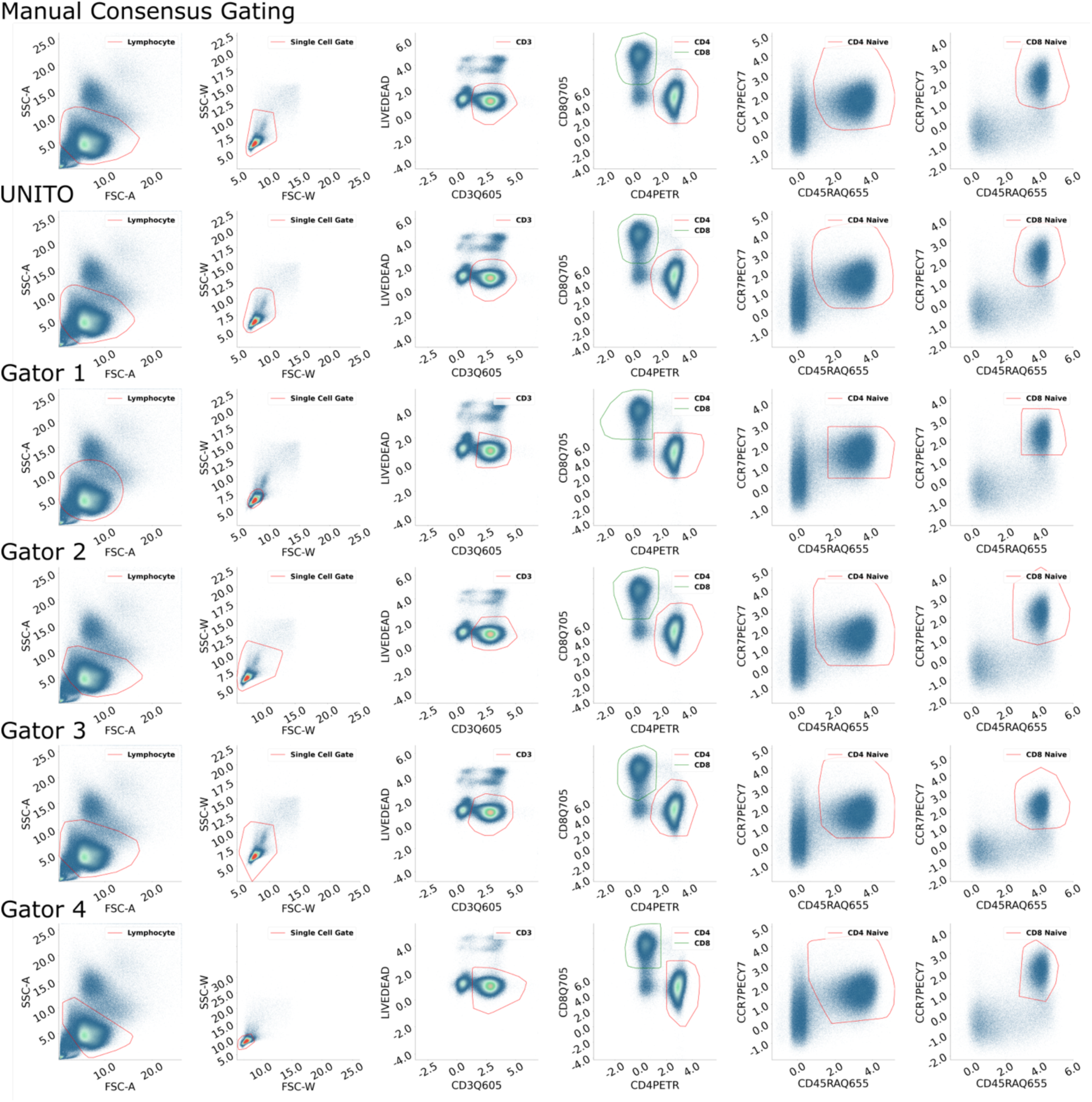
Gating results by gaters (flow cytometry). With the same aim to assess the performance of UNITO compared to manual gaters, we plotted the same format using the flow cytometry dataset (top to bottom: consensus manual gating, UNITO gating, gater 1-4). The gates are organized in the same way as previous figures (left to right: lymphocyte, single cell gate, CD3, CD4 + CD8, CD4naive, CD8naive).

**Figure S6.**
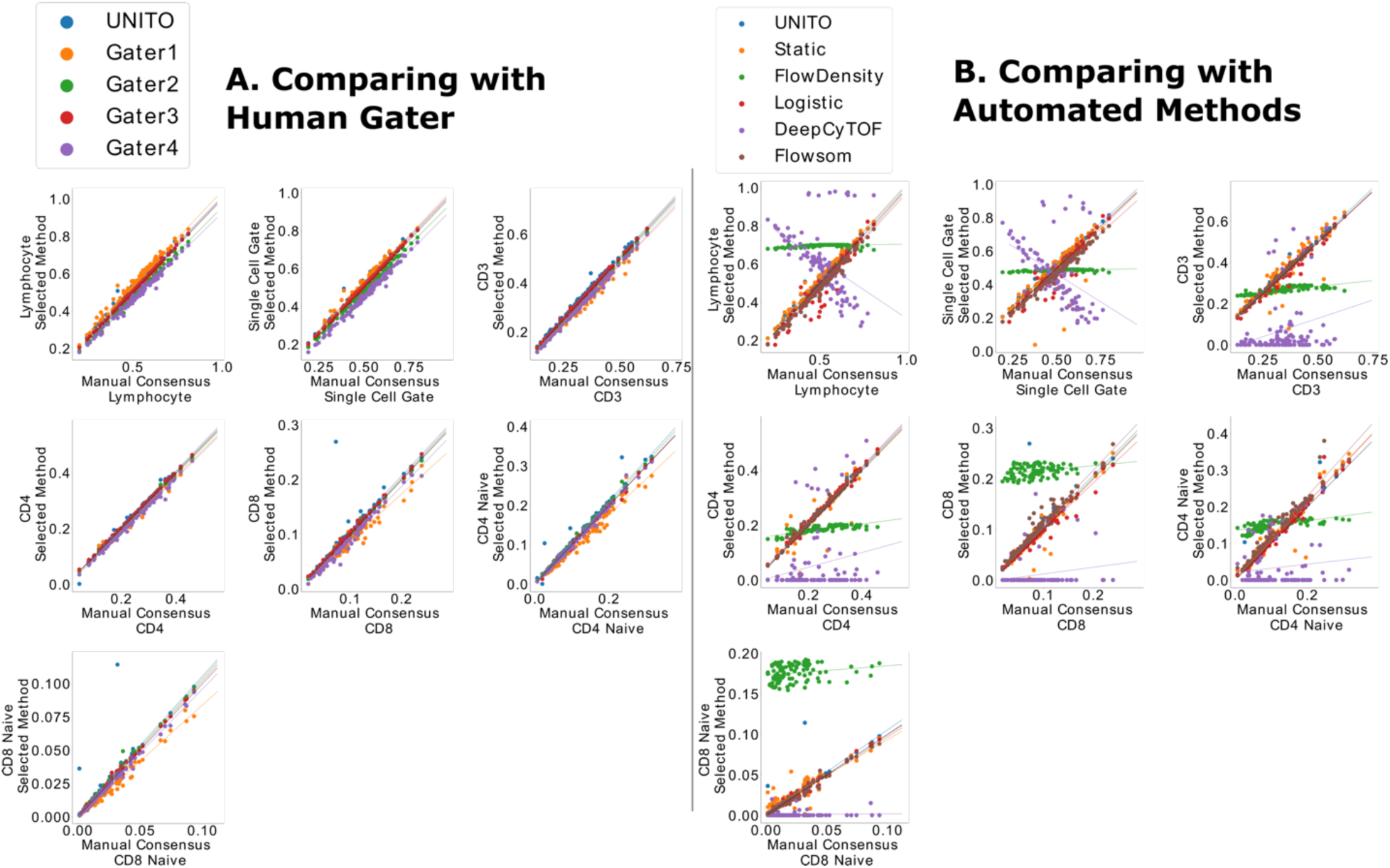
Proportion of cells over the entire cell population captured by manual gating versus human/automated gating in flow cytometry (left for vs human and right for vs automated methods). Each dot in the plot represents a single subject. The plot is separated by every single gate, and within the same coordinate results from each method were visualized by different colors. The dashed line represents a perfect correlation with manual consensus gating, and the Pearson correlation coefficients are reported in the supplementary material.

**Figure S7.**
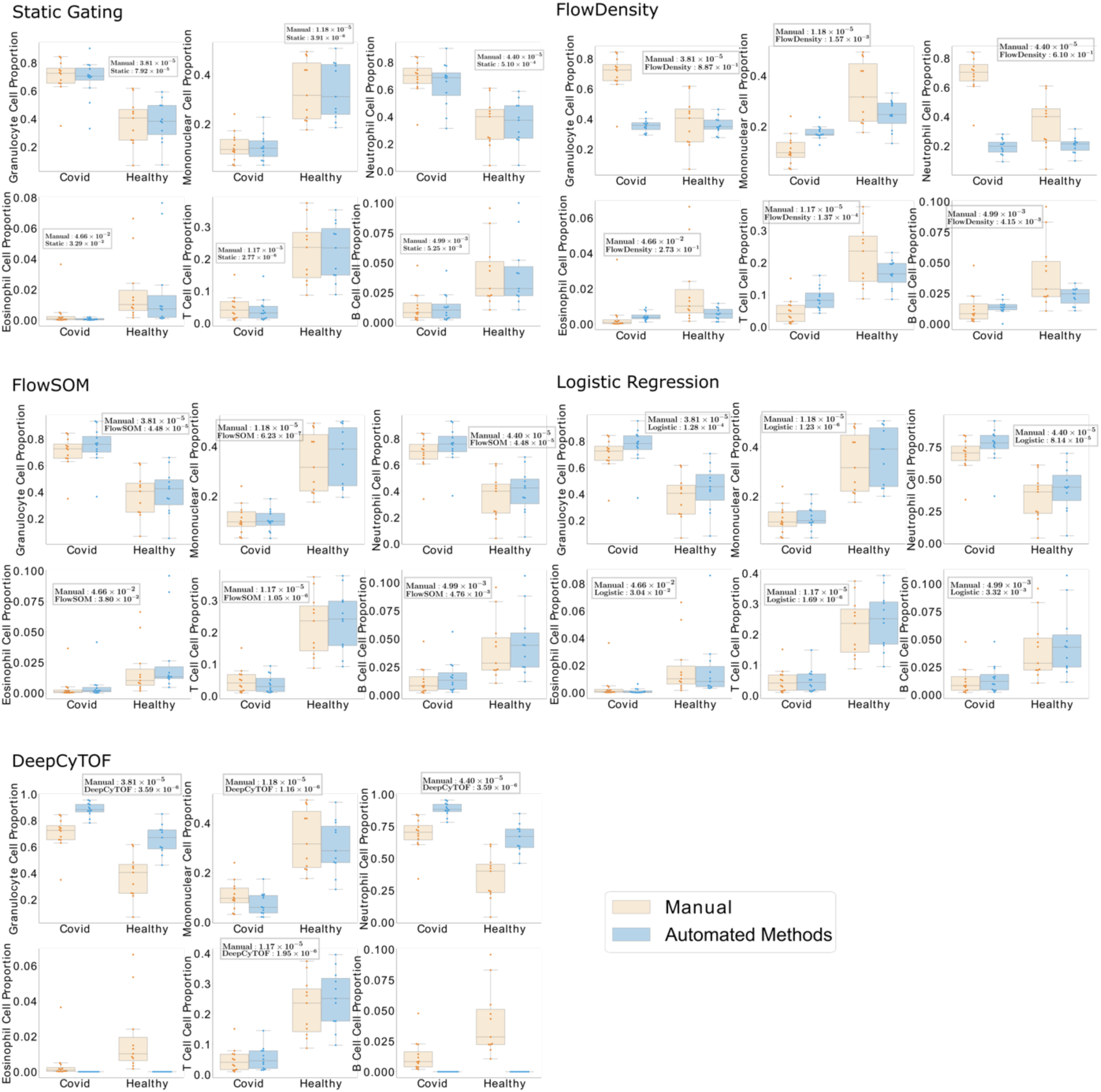
Statistical analysis on group differences between COVID-19 subjects and healthy subjects (mass cytometry). Downstream analysis shows the methods’ ability to reveal further biological insights after gating. Here we visualized the group-level comparison for selected cell types in mass cytometry gating from all comparison methods (top left: static gating, top right: FlowDensity, middle left: FlowSOM, middle right: logistic regression, bottom left: DeepCyTOF). Within each subplot, we selected 6 intermediate and bottom level gates in the gating hierarchy (top row from left to right: granulocyte, mononuclear, neutrophil; bottom row from left to right: eosinophil, t cell, and b cell). Box plots and swarm plots of manual gating and automated gating are plotted together for easier comparison between them, and the group-difference p-value by t-test is reported for each panel.

**Figure S8.**
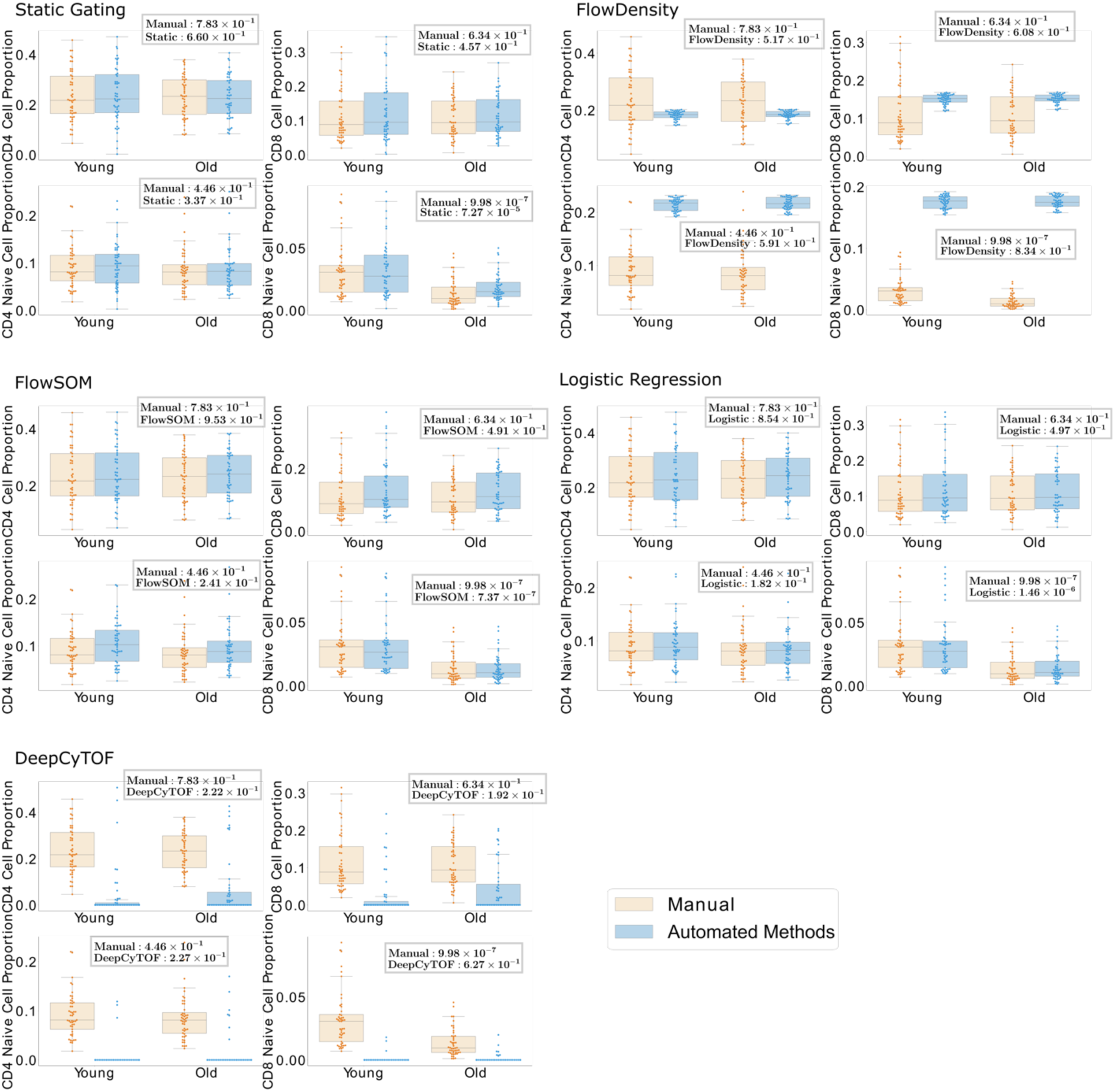
Statistical analysis on group differences between young and old subjects (flow cytometry). The same downstream analysis was applied to the flow cytometry dataset to distinguish young and old subjects. The methods were organized in the same order as the Figure S7, and four cell types were selected for downstream evaluation (top left: CD4, top right: CD8, bottom left: CD4 naïve, bottom right: CD8 naïve). The same group with both manual and automated methods are plotted together to see the disparity, and p-values from the t-test between young and old groups are reported.

**Figure S9.**
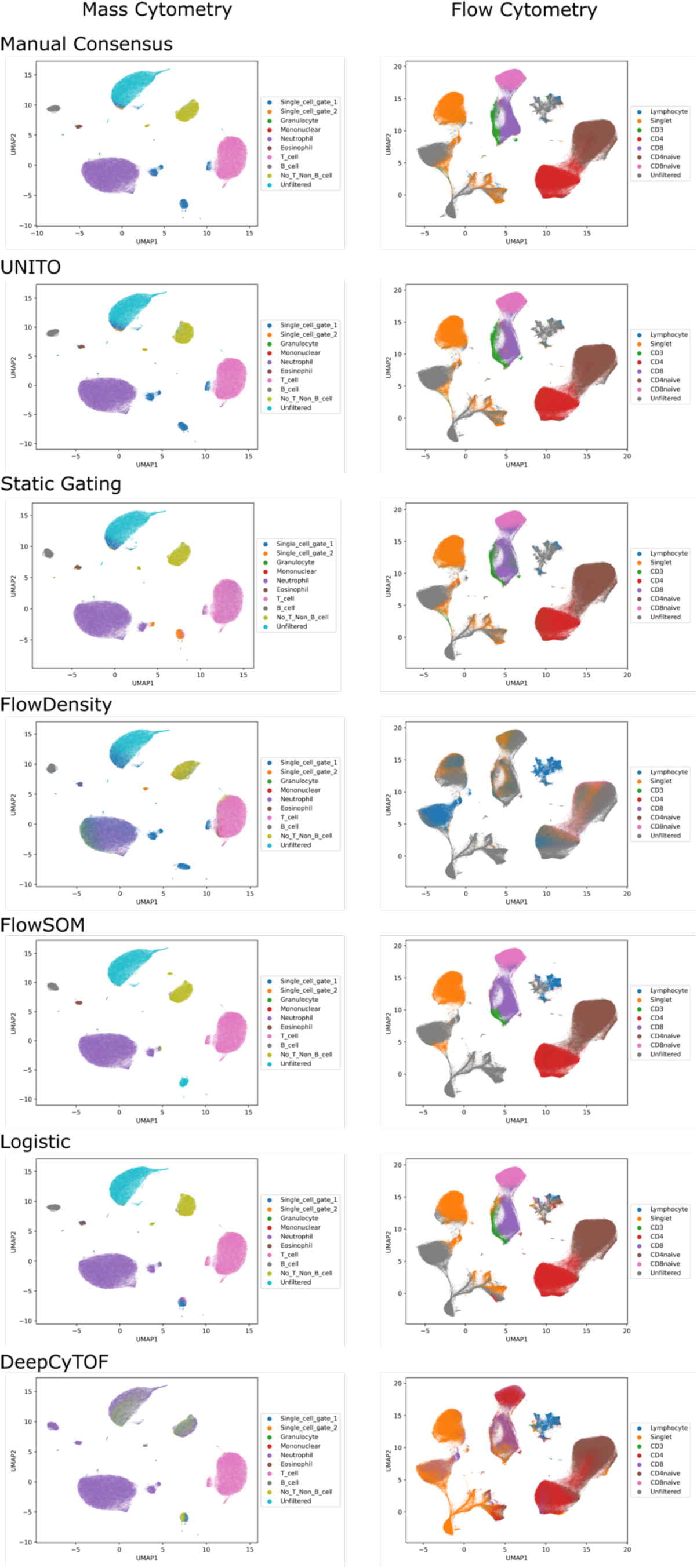
UMAP clustering after gating. UMAP was applied to the gating results for each method to see the separation between different cell types (top to bottom: manual consensus, UNITO, static gating, FlowDensity, FlowSOM, logistic regression, DeepCyTOF). Since we have a hierarchical gating structure, the cell type label corresponds to the higher level only contains the cell that are belong to the current gates but have been excluded from subsequent gates (for example, the cells from single cell gate 1 only contain those cell that are classified as single cell gating 1 but at the same time not identified as single cell gate 2).

